# Ancient herpes simplex 1 genomes reveal recent viral structure in Eurasia

**DOI:** 10.1101/2022.01.19.476912

**Authors:** Meriam Guellil, Lucy van Dorp, Sarah A. Inskip, Jenna M. Dittmar, Lehti Saag, Kristiina Tambets, Ruoyun Hui, Alice Rose, Eugenia D’Atanasio, Aivar Kriiska, Liivi Varul, A.M.H.C. Koekkelkoren, Rimma D. Goldina, Craig Cessford, Anu Solnik, Mait Metspalu, Johannes Krause, Alexander Herbig, John E. Robb, Charlotte J. Houldcroft, Christiana L. Scheib

## Abstract

Human herpes simplex virus 1 (HSV-1), a life-long infection spread by oral contact, today infects a majority of adults globally^1^, yet no ancient HSV-1 genomes have yet been published. Phylogeographic clustering of sampled diversity into European, pan-Eurasian, and African groups^2, 3^ has suggested that the virus co-diverged with anatomically modern humans migrating out of Africa^4^, although a much younger origin has also been proposed^5^. The lack of ancient HSV-1 genomes, high rates of recombination, and high mobility of humans in the modern era have impeded the understanding of HSV-1’s evolutionary history. Here we present three full ancient European HSV-1 genomes and one partial genome, dating to between the 3rd and 17th century CE, sequenced to up to 9.5× with paired human genomes up to 10.16×. These HSV-1 strains fall within modern Eurasian diversity. We estimate a mean mutation rate of 7.6 × 10^-7^ - 1.13 × 10^-6^ for non-African diversity leading to an estimated age of sampled modern Eurasian diversity to 4.68 (3.87 - 5.65) kya. Extrapolation of these rates indicate the age of sampled HSV-1 to 5.29 (4.60-6.12 kya, suggesting lineage replacement coinciding with late Neolithisation and implicating Bronze Age migrations^6^ in the distribution of HSV-1 through Eurasia.

## Main

Humans are hosts to a large number of viruses. For many, major uncertainties exist regarding how long they have infected humans, how associated disease pathologies may have changed since their emergence, and the extent to which patterns of viral diversification may mirror the interactions and behaviours of their hosts. Co-analysis of host and pathogen genomes offer rich opportunities to address these questions; however, such studies are often challenged by genomic sampling spanning over relatively recent time-scales. When relying on modern data, important assumptions have been made regarding pathogen mutation rates and joint demographic histories, only to be overturned by ancient genomic data^7–9^. Herpes viruses are a prime example of human pathogens for which assumptions have been necessary due to the lack of ancient genomes.

Herpes simplex virus 1 (HSV-1) is a double-stranded DNA virus that affects at least two-thirds of the human population. The majority of people are infected in infancy or childhood. Most infections are mild or asymptomatic; however, HSV-1 can, in rare cases, lead to severe complications especially in those immuno-compromised due to underlying genetic predisposition, infection or malnutrition^1^. Following primary infection, the virus becomes latent in sensory neurons^10^. When triggered by psychological or physiological stress, the virus can reactivate resulting in recurrent labial lesions^11, 12^. The emergence of neutralising antibodies and cytotoxic T cell responses ensure reactivations typically do not lead to substantial viraemia in the immunocompetent host^13^. A number of human genetic variants have been implicated in susceptibility to HSV-1 in otherwise healthy individuals, covering a range of phenotypes from mild^14, 15^ to severe, such as the association between TLR3 deficiency and fatal HSV-1 encephalitis^16^.

Despite the high global prevalence of HSV-1, there are relatively few genomes available, with modern diversity clearly undersampled^17^. Comparative studies have indicated that HSV-1 can be clustered into three ‘phylogroups’ associated with geographic origin: I (Europe and America), II (Europe, Asia and America), and III (Africa) ^2, 18, 19^. Such a distribution, with phylogroup III falling basal to Eurasian phylogroups, has led to speculation that the origin and subsequent apportionment of genomic diversity in HSV-1 coincided in tandem with human migrations out of Africa^4^. This co-divergence scenario has also been suggested for other agents of common childhood infections^20–22^. Conversely, recent work has suggested that the origin of HSV-1 is far more recent, with little geographic clustering outside of African strains and an estimated time to the most recent common ancestor (TMRCA) dating to less than 7,000 years ago^5^. To date, the majority of modern HSV-1 genomes are partial (85%) and/or clinical isolates from highly cosmopolitan centres^18, 23–26^. The oldest HSV-1 genome (strain HF) was isolated from an individual living in New York in 1925^27^ and like many laboratory strains has been passaged for generations before being sequenced, which can lead to higher rates of evolution than in natural conditions. Furthermore, recombining viruses, such as HSV-1, may undergo high rates of lineage replacement; meaning diversity sampled over recent shallow time depths offers limited power to resolve more ancient origins^28^ while also leading to faster estimates of evolution^29^. Without direct ancient calibration points, it is difficult to assess the processes giving rise to modern sampled diversity.

Ancient DNA (aDNA) has become an increasingly powerful tool for studying past infections, as massive parallel sequencing (NGS) has allowed for DNA libraries extracted from skeletal samples to be screened for thousands of microbial species (reviewed in Syprou et al. 2019^30^), shedding light on the evolutionary history and phylogeography of past infections. HSV-1 DNA is detectable in blood during a primary infection, but not during reactivation from latency^31^; however, herpesvirus DNA and microRNA from non-primary infections have been recovered from the teeth^32^ and subgingival plaque^33–35^ of living individuals. HSV-1 DNA has also been recovered from the trigeminal ganglia of cadavers^12^ indicating that it can reactivate peri- or post-mortem. HSV-1 should therefore be common in the archaeological record; however, to date no full ancient HSV-1 genomes have been published.

### Recovery of ancient HSV-1 genomes

We identified four aDNA libraries generated from teeth that contained HSV-1-specific reads. The samples came from a young adult male from an urban medieval hospital cemetery (JDS005; 1350-1450 CE) and an adult female from an early Anglo-Saxon cemetery (EDI111; 500-575 CE) in Cambridgeshire, United Kingdom; an adult male from a burial related to the Nevolino culture (BRO001; 253 - 530 CE) in Russia; and an adult male from the Netherlands (RIJ001; 17th century) (Extended Data Table 1, Supplementary Note 1). These libraries were then sequenced to higher depth with additional shotgun sequencing or target enrichment to achieve paired human and viral ancient genomes at 0.03 - 11× and 1.2 - 9.5× average coverage respectively (Fig. 1A, Supplementary Table 1A,B). In both human and HSV-1 genomes, terminal misincorporation patterns are consistent with aDNA postmortem damage (Fig. 1B, Supplementary Table 1A,B) and estimated human contamination rates are low (Extended Data Table 2). The RIJ001 alignment was only partial and showed low depth of coverage across the entire reference sequence (Fig. 1A), we therefore excluded the sample from most analysis if not otherwise specified.

**Fig. 1.**
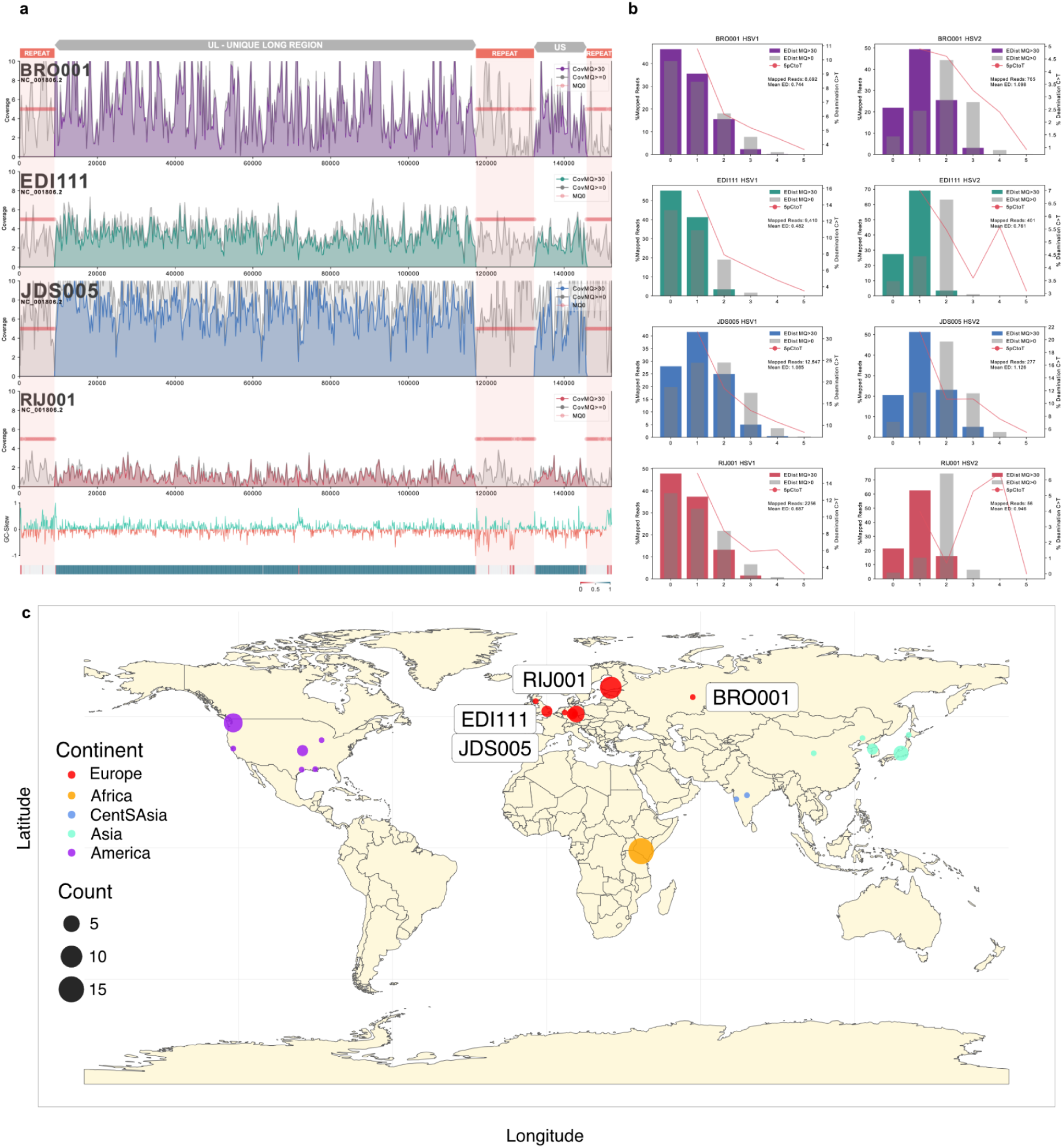
Sequence coverage and edit distance of samples analysed. **a,** Linear plots of our mappings to the reference strain 17. The first four plots represent sequence coverage (150bp windows) and depth across the reference strain for each sample. The last two plots show the GC-skew and the mappability estimates based on GenMap^55^ (<0.4 in red; >0.6 in blue) . **b,** Edit distances for genomes mapped against HSV-1 and HSV-2. Note JDS005 exhibits more postmortem damage than other genomes and is more divergent from the reference sequence, leading to higher edit distance from HSV-1. **c,** World map providing the sampling location of modern HSV-1 accessions jointly analysed with three ancient samples: EDI111, JDS005 and BRO001 (as highlighted). Included accessions are provided in Supplementary Table 5. Colour provides the continent of sampling and size the sample count per location as per the legend at left.

Given the scarcity of HSV-1 in the ancient genomics record we assessed whether these genomes were retrievable due to increased susceptibility of the individuals to infection. We tested for known susceptibility variants or human leukocyte antigen (HL) alleles implicated in susceptibility to fatal HSV-1, or high runs of homozygosity (ROH), which can lead to haploinsufficiency (Supplementary Note 6) in the samples with > 1X autosomal coverage. Host genotypes were called in JDS005 and EDI111; however, neither of these individuals carried variants suggested to cause rare, deadly susceptibility to herpes infections (Fig. 2A, Supplementary Table 2) nor runs of homozygosity extending over 4cM (Extended Data Table 3). We called HLA A, B, and C alleles in both individuals (Fig. 2B) using software designed for next-generation sequencing data^36^. Notably, only one HLA-B haplotype was detected in EDI111. Isotope data from tooth dentine and bone collagen of JDS005 indicates a diet comparatively low in animal or marine proteins, perhaps due to poverty and/or an inability to take in sufficient nutrition due to other health conditions (Supplementary Note 7). No isotopic information is available for the other three individuals.

**Fig. 2.**
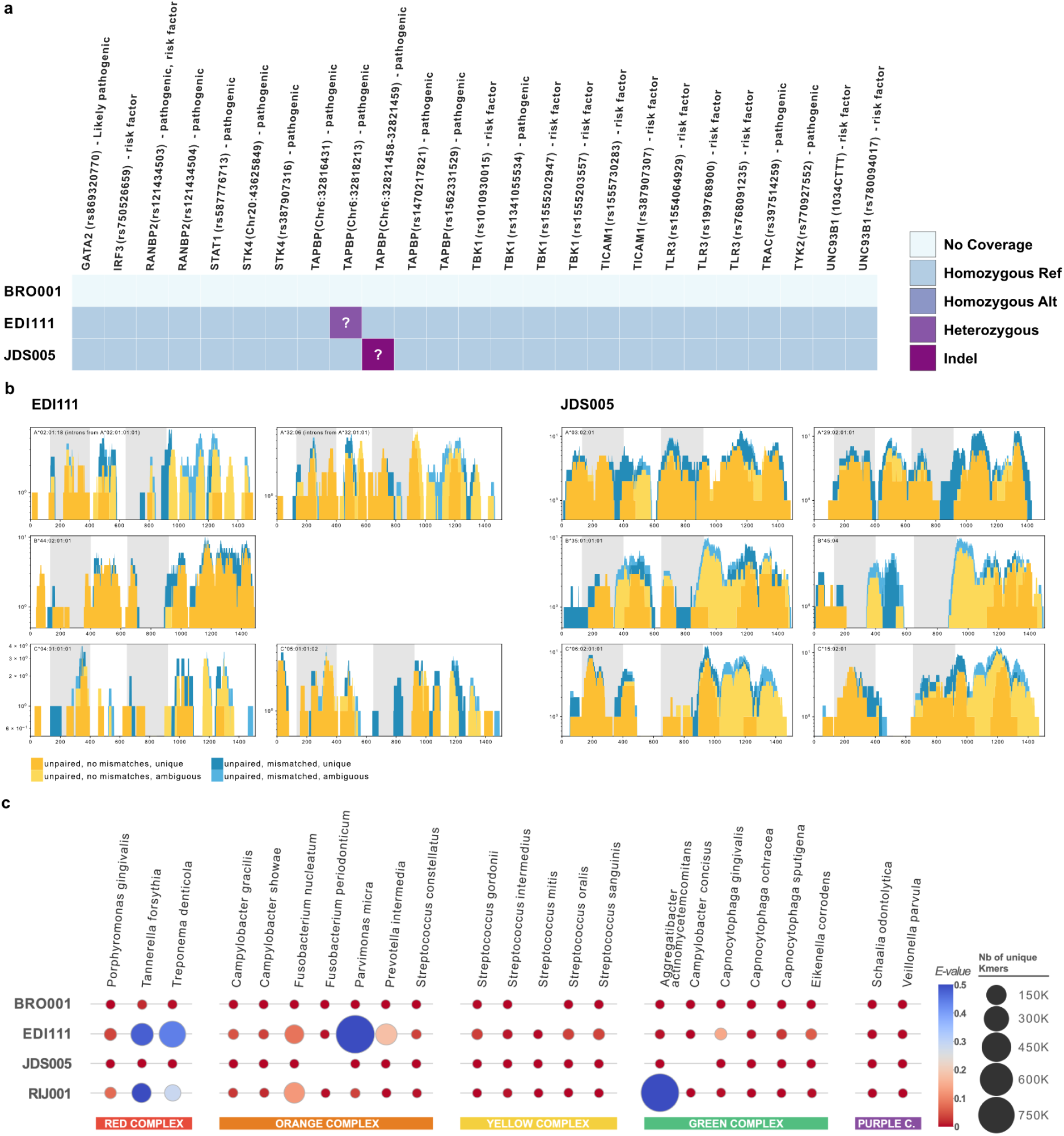
Host susceptibility factors. **a,** Reference vs. alternative alleles in the host genomes of mutations in genes related to susceptibility to HSV infections as categorised by the ClinVar database downloaded on 17/06/2021. See Supplementary Table 2 for more information. **b,** Coverage and haplotype assignment of HLA-A, HLA-B, and HLA-C alleles in the two individuals with human genomic coverage over 3x. Graphs were generated using OptiType^36^. **c,** Heatmap showing the unique k-mer hits (size) and E-value (hue) calculated (see methods) for all four samples based on KrakenUniq data. Represented are species associated with periodontal disease grouped by complexes based on Socransky et al. 1998 ^39^ for which hits could be identified for at least one sample.

We then assessed whether the three full ancient HSV-1 strains exhibited genotypic markers of predicted high consequence mutations/variations which may lead to higher pathogenicity in the host. Based on our SNP effect analysis, SNPs within intervals associated with pathogenicity, immunogenicity or known cellular phenotypes were investigated (see SI note 5). Of these, the strain BRO001 sampled in Russia carried a mutation leading to a stop-loss in the *US3* domain; however, it is unclear whether this would have affected transcription^37^. This strain also carried a number of SNPs inferred to have a moderate phenotypic impact in the *UL22/gH* domain (the viral envelope protein), increasing the likelihood that this variation may encompass an antigenic region and thus may be recognised as B or T cell epitopes. No known phenotypically relevant mutations were identified in the Cambridge strains (Supplementary Note 5).

HSV-1 has been found at higher prevalence in cases of chronic and aggressive periodontitis^34, 35, 38^, though the role in pathogenesis is inconclusive^38^. Both EDI111 and JDS005 have dental pathology consistent with periodontal disease (Supplementary Note 1). Metagenomic screening identified the presence of sequences commonly associated with periodontal pathogens in EDI111 and RIJ001 (Fig. 2C, Supplementary Table 4) including *Tannerella forsythia, Treponema denticola, Fusobacterium nucleatum*, as well as *Parvimonas micra* and *Aggregatibacter actinomycetemcomitans* for EDI111 and RIJ001 respectively.

*Tannerella forsythia* and *Treponema denticola* form part of the so-called “red-complex,” which has been strongly implicated in periodontal disease^39^. While these results cannot verify the presence of periodontitis in EDI111 and RIJ001, the presence of a range of these bacteria in sufficient amounts to be easily detectable points towards the presence of an infection.

The sum of available evidence indicates that the ancient HSV-1 infections we retrieved were typical, possibly recurrent infections, though these individuals could have had immuno-compromising complications just prior to death leading to an increased viral load.

### Evolutionary history of HSV-1

To place our full ancient HSV-1 genomes into extant diversity, we curated a dataset of modern genome assemblies spanning Europe (United Kingdom, Germany, Finland), Central and East Asia (China, South Korea, Japan and India), Africa (Kenya) and the Americas (USA) (Supplementary Table 5, Figure 1C). While additional genomes are available from the USA, we subsequently restricted our analysis to those sampled prior to 1989 to limit the inclusion of highly cosmopolitan strains in our dataset and to maximise geographic structure. We combined the modern dataset with the three ancient higher coverage genomes using a core SNP calling approach and masked hyper diverse and repeat regions.

A phylogenetic network was created over the core alignment of modern and ancient HSV-1 using SplitsTree4^40^ (Fig. 3A). The median spanning network recovered major stratification of phylogroups I, II and III though with high levels of recombination underpinning the clustering. Of note, each of the ancient genomes falls within the diversity observed in Eurasian HSV-1 clusters; broadly defined by phylogroup I and II^2^, a pattern consistent regardless of the reference genome used (Extended Data Fig. 1). Given the excess of recombination identified in internal nodes of the network we formally tested for the presence of recombination in the core alignment; detecting a significant correlation between linkage disequilibrium over genetic distance (*p*=5.73e-8) (Extended Data Fig. 2A). We therefore proceeded to employ population genetics methods to identify genetic clusters within our dataset and to characterise the affinity between modern and ancient HSV-1 strains.

**Fig. 3.**
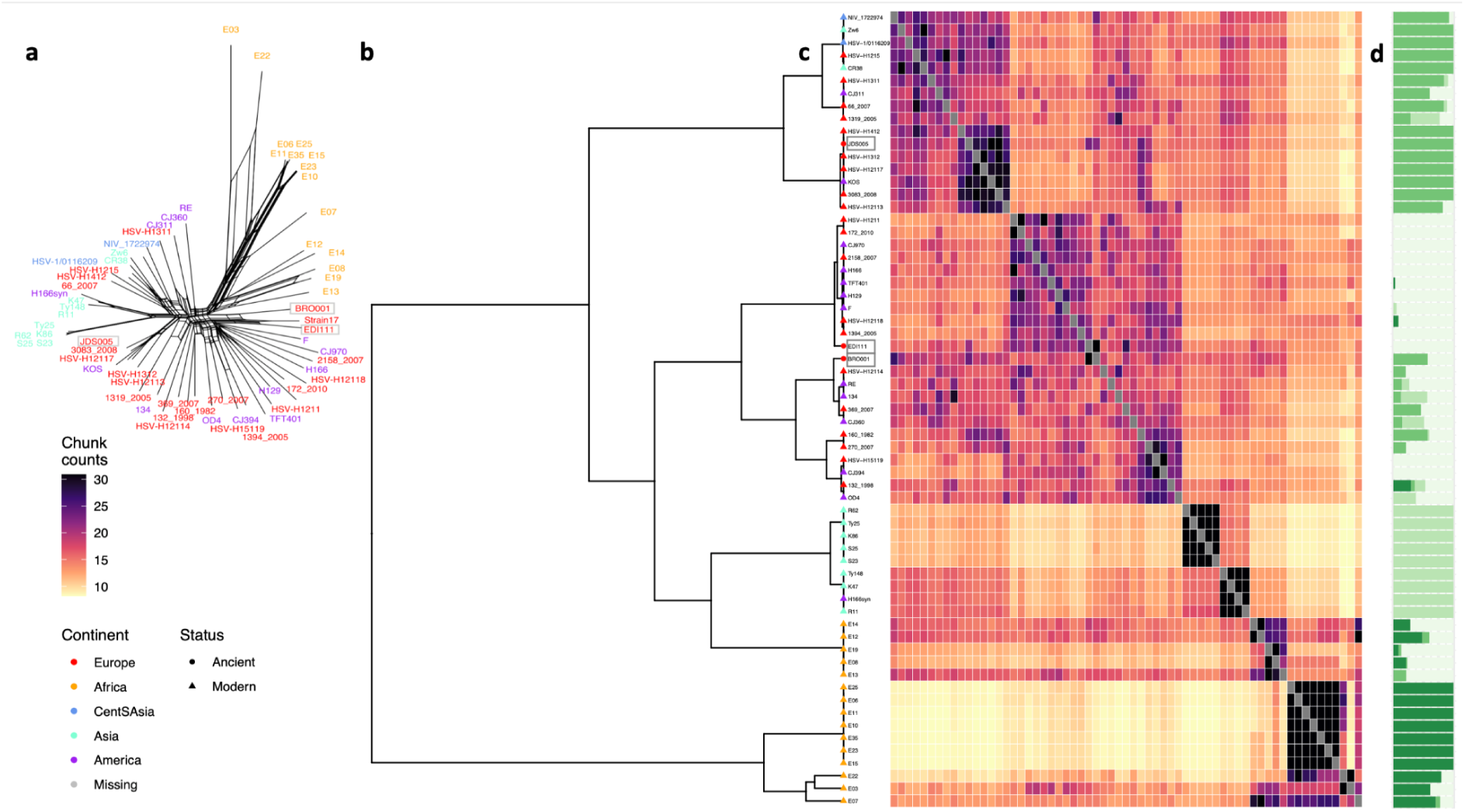
Phylogenetic distribution of HSV-1 genomes. **a,** SplitsTree neighbour net network of three ancient HSV-1 (highlighted by grey boxes) and curated set of modern global genomes^2, 18, 19, 56–58^. Label colours correspond to the continent of origin (legend at bottom left). **b,** fineSTRUCTURE hierarchical clustering of all HSV-1 over the core genome alignment. Tips are coloured by continent of sampling and ancient/ modern status as given by the legend at bottom left. **c,** Heatmap providing the average number of haplotype chunks shared between any two considered genomes, with colour scale given at bottom left. **d,** Unsupervised clustering analysis implemented in ADMIXTURE at K=4 ordered as per fineSTRUCTURE inferred hierarchical clustering.

Unsupervised model-based clustering^41^ recovered four well-supported clusters (Extended Data Fig. 2B), again largely delineating the tripartite diversity separating African HSV-1 from European-like ‘phylogroup I’ and Eurasian ‘phylogroup II’; with our two oldest high coverage genomes sharing inferred ancestry components dominant in European and American strains; and with the more recent ancient HSV-1 genome (1350-1450 CE) enriched for an ancestry component identified largely in ‘phylogroup II’ (Fig. 3C). To further resolve patterns of genetic diversity we applied a haplotype-based clustering method^42^, utilising chromosome painting. Highly consistently, patterns of haplotype sharing partitioned three major groupings with ancient European HSV-1 falling within the two major extant Eurasian clades (Fig. 3B,C).

As some modern HSV-2 strains have recombined with HSV-1^43, 44^, there is a possibility that HSV-1 strains may in turn carry HSV-2 x HSV-1 recombinant regions; however, when assessed using RPD5^45^, we found no evidence of recombination with HSV-2 in our ancient strains. We next pruned our alignment for the presence of putative recombinant tracts within HSV-1 strains (Extended Data Fig. 2C, Supplementary Table 6) as their inclusion may violate assumptions of standard phylogenetic approaches. Using the alignment filtered for detectable recombination we constructed a maximum likelihood phylogeny (Extended Data Fig. 3). The phylogeny largely recapitulated results from recombination-aware analyses, though resolved notable sub-structure within African HSV-1 sampled in Nairobi^19^, an observation consistent with an African origin. The two oldest ancient genomes fall basal to the diversity of phylogroup I strains, while our medieval genome is placed in phylogroup II, closest genetically to modern strains sampled in Germany and Finland.

We assessed whether the extant diversity observed in HSV-1 could be best explained by a deep co-divergence with human hosts, or instead derives from more recent prehistory. To do so, we tested for the significant accumulation of mutations over the time of sampling in our dataset; taking forward those genomes with reported sample collection dates (Supplementary Table 5). While we were unable to recover a robust global temporal signal; we detected a significant correlation, robust after date randomisation resampling, at the node ancestral to all Eurasian samples including our ancient genomes (R2=0.2; *p*=0.006) (Extended Data Fig. 4A,B). Using the time of sample collection and radiocarbon estimates for our ancient samples as priors, we estimate the substitution rate and most recent common ancestor of Eurasian diversity, testing six different demographic models pre-specifying relaxed and strict priors on the evolutionary rates^46^. Models converged to highly similar estimates, with posteriors significantly different from those estimated when sampling from the prior. This resulted in estimated rates of 2.38 × 10^-7^ - 2.49 × 10^-6^ substitutions per site per year (Extended Data Fig. 4A) and a time to the most recent common ancestor of sampled Eurasian HSV-1 strains spanning 1.87 - 12.73 kya across models, with our highest likelihood implementation supporting a MRCA of 4.67 (3.78 - 5.65) kya (Fig. 3A, Extended Data Table 4, Extended Data Fig. 4C,D). We estimate the common ancestor of BRO001 and EDI111 to be 3.45 (2.99-3.95) kya; with the estimated MRCA of European phylogroup I and pan-Eurasian phylogroup II to approximately the same time periods of 4.50 (3.77-5.30) kya and 4.47 (3.64-5.33) kya respectively overlapping the Early Bronze Age in Europe (Fig. 3B). Two Asian monophyletic clades were placed within the diversity of phylogroup II, dating to 1.32 (1.50-1.63) kya and 123.05 ya (89.67 ya - 161.70ya) respectively. Fixing these estimated rates over the global phylogeny, which includes a further 15 African HSV-1 strains, results in an estimate of extant circulating diversity to 5.29 (4.60-6.12) kya (95% HPD) (Extended Data Fig. 4B).

## Discussion

Given the high prevalence of HSV-1 infections in human populations today and the pathophysiology of the virus, HSV-1 should be abundant in archaeological teeth. Our study targeted the apical root and the proximity to neurons may have facilitated HSV-1 detection^47^. HSV1 is typically found in both trigeminal ganglia (TG) of infected individuals, but with limited genome replication during latent infection^47^. Inter-individual variability in the proximity of apical roots to the nerve canal may be relevant here too^48^. One possibility for the lack of ancient HSV-1 genomes observed to date is that most individuals have a latent infection with low HSV-1 copy numbers in the TG or reduced oral shedding due to a robust immune response^49^. Today, Herpes viruses are more likely to appear in the presence of periodontal disease or other inflammation^34, 35^, thus HSV-1 might be more likely to be found in ancient individuals suffering the same. Factors that lead to higher oral shedding should be investigated to better inform sampling strategy for ancient HSV-1 studies.

Since we do not detect large-scale changes in the core genomic composition of ancient and modern HSV-1 strains (Supplementary Table 7), one possibility is that HSV-1 may have become more prevalent over time due to changes in host behaviour or altered viral transmission routes. Changing transmission dynamics may offer an explanation to either a reasonably recent emergence of extant HSV-1 or lineage replacement in the history of Eurasian HSV-1, as implicated by the relatively recent dates we obtain. The primary mode of HSV-1 transmission is vertical, from parent to child; however, the addition of lateral transmission as population density increased during the Bronze Age, potentially linked to cultural practices such as the advent of sexual-romantic kissing, may have contributed to a shift in the dominant lineages which have continued to circulate to this day.

A more recent origin of sampled Eurasian HSV-1 genotypes has support from other studies of modern data^5^ and in alphaherpesviruses^50^. For instance, Weinert and colleagues dated the emergence of circulating varicella-zoster virus diversity to within the last 5,000 years, and used rates of live attenuated vaccine evolution to suggest that an Out of Africa scenario was implausible^50^. While VZV is spread by infectious aerosols, its substitution rate is expected to be similar to that of other *Alphaherpesvirinae*^29^. *Mycobacterium tuberculosis* was also thought to have much older origins and shared co-evolutionary history with humans^51^; however, analyses relaxing the assumptions of co-divergence point to a much more recent history^52^. Similar approaches have been used to estimate conflicting ages of HSV-1^4, 5^ This does not preclude an ancient association of HSV-1 with human hosts. For instance, recent work suggests the presence of putative Herpesvirus reads in children dating to 31kya^22^. Nonetheless our results, uniquely aided by observations from full ancient genomes, suggest that the distribution of extant HSV-1 is the product of more recent events.

The four ancient HSV-1 strains we recover also aid elucidation of the origins of modern strains, for example, the placement of the KOS strain close to the medieval Cambridge and other European sequences suggests Europe as a more likely origin of this strain than Asia as was hypothesised in earlier work^53, 54^, nor is it likely to be an indigenous American strain^4^. However, even with the inclusion of ancient genomes, the highly cosmopolitan nature of modern sampling is not ideal for reconstructing historic HSV-1 transmission events, with all estimated TMRCAs highly sampling dependent. Our work therefore highlights the need for more extensive coverage of modern HSV-1, particularly in regions such as Asia and Africa, together with additional observations provided by aDNA samples. Further ancient genomes, for example from the Neolithic period, may further revise our understanding of the evolutionary history of this today ubiquitous pathogen and continue to inform on the nature of its association with human hosts.

## Methods

No statistical methods were used to predetermine sample size. The experiments were not randomized and investigators were not blinded to allocation during experiments and outcome assessment.

### Ethics statement

All skeletal elements were sampled with permissions from the representative bodies/host institutions. Samples were taken and processed to maximise research value and minimise destructive sampling.

### Selection of archaeological samples

The individuals included in the study were sampled as part of larger interdisciplinary or population genetics studies and thus were not targeted for ancient viral analysis. Teeth were sampled from skeletons while wearing gloves. Molars were preferred due to higher mass and in this case proximity to the trigeminal nerve may have been a factor in discoverability of HSV-1 DNA.

### Generation of ancient DNA sequence data

Sampling, decontamination, extraction, purification and library preparation for JDS005 was carried out as described in Scheib et al. 2018^59^. EDI111 and RIJ001 were processed at the Institute of Genomics ancient DNA facility at the University of Tartu as described in Scheib et al. 2019^60^. BRO001 was processed at the Institute of Ecology and Earth Sciences ancient DNA facility at the University of Tartu as described in Saag et al. 2019^61^.

### Radiocarbon determinations

A radiocarbon determination was generated for EDI11 from the tooth at the ^14^Chrono Centre, Queen’s University Belfast (Supplementary Table 8) and for BRO001 at the Poznan Radiocarbon Laboratory under the ID Poz-98180.

### Metagenomic screening

Generated shotgun sequencing data was inspected with fastqc^62^, and quality filtered and trimmed of adaptors using cutadapt^63^ with the settings -m 30 --nextseq-trim=20 for single-end data with multi-core support enabled (-m 30 --nextseq-trim=20 --times 3 -e 0.2 -j 0 --trim-n). We then deduplicated the filtered FASTQ files using ParDre^64^. Following these steps, microbial DNA was assigned using KrakenUniq^65^against a custom database of complete genomes and chromosome level assemblies of bacteria, viruses, archaea and protozoa. The human genome, the NCBI Viral Neighbor database and the contaminant databases UniVec and EmVec were also included. A custom E-value calculated as follows: 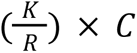. Here, K is the *k*mer count, R is the read count and C is the coverage of the taxon *k*mer dictionary. The E-value cut-off for further inspection was 0.001.

### Target capture of HSV-1

Due to low coverage, we enriched the shotgun library of sample BRO001 and RIJ001 for HSV-1 DNA using an Arbor Biosciences Custom MyBaits multispecies viral capture kit (v4), that includes sequences from 283 *Herpes simplex virus type 1* genomes and 261 *Herpes simplex virus type 2* genomes. The captured library was amplified using 2X KAPA HiFi HotStart ReadyMix DNA Polymerase and primers IS5 and IS6^66^. Following amplification, the library was sequenced on a NextSeq500 platform (150bp, paired-end) at the Core Facility of Institute of Genomics.

### Mapping and authentication of viral data

Raw data was returned in the form of single and paired FASTQ files. Prior to mapping single-end datasets were trimmed and filtered as described for the metagenomic screening. Paired-end datasets were also filtered and trimmed using cutadapt^63^ (-m 30 --nextseq-trim=20 --times 3 -e 0.2 -j 0 --trim-n --pair-filter=any). Forward and reverse reads were merged using FLASH^67^ (-M 125).

All datasets were merged by library and mapped non-competitively against the *Herpes simplex type 1* reference sequences for strain 17 (<u>NC_001806.2</u>), strain S25 (<u>HM585513.2</u>), strain 2158_2007 (<u>LT594106.1</u>) and strain E07 (<u>HM585497.2</u>), to exclude a reference bias, using bwa aln (-n 0.1 -l 1000) and bwa samse^68^. We then used samtools^69^ to convert and sort the data to bam format. Duplicates were marked and removed using Picards’s^70^ MarkDuplicates module. We then merged sequencing runs for all libraries using samtools merge (-R) and realigned reads around indels using GATK’s^71^ RealignerTargetCreator and IndelRealigner modules. We estimated deamination rates using mapDamage2.0^72^. Alignments were visualized using circos^73^ and seaborn^74^ and mapping statistics were collected using Qualimap2^75^. Edit distances were computed with the ED-NM_CSVscript^76^ using the modules numpy^77^, pysam^78^ and pysamstats^79^. Additionally, we also mapped our data non-competitively to the *Herpes simplex type 2* strain HG52 reference genome (<u>NC_001798.2</u>) using the same workflow.

All positions were stored using samtools v1.10 mpileup (*-aa* flag)^69^. Variant positions were then called using bcftools v1.10.2, filtering for sites with a mapping quality of at least 20, a depth of 3, a quality of 20 and taking forward those positions with an alternate allele fraction >0.8 (MQ>20, DP>3, Q>20, AAF>0.8). Consensus fasta files were generated using bcftools *consensus* masking those regions of the genome not covered over the reference. Variants were called for all included modern genomes from their respective assemblies using the snippy pipeline and specifying contigs as input (https://github.com/tseemann/snippy). The resulting ancient and modern variants were then considered to jointly construct a core-genome alignment, as in Lassalle et al^80^, using snippy-core, taking forward positions observed in at least ten accessions across the dataset and masking difficult to call positions including repeat genes (*RL1, RL2, RS1*) together with all regions identified by Szpara et al. 2014^19^. Masking resulted in exclusion of 12,614 nucleotides of the reference (8%) and a resulting SNP alignment of 3,815 well defined variant positions distributed evenly over the remaining genome across 63 accessions (Extended Data Figure 5). The distribution of SNPs per accession was 281-669 (95% CI) with each of the higher coverage ancient genomes (BRO001, EDI111, JDS005) having 275, 185 and 286 core SNPs respectively. Equivalent analyses were conducted using three representative alternate references to assess the placement of the ancient strains.

We estimated the mappability across the reference sequence strain 17 using GenMap^55^ (-K30 -E2). Coverage across gene intervals was calculated using bedtools coverage^81^ and mean mappability across these intervals was computed using bedtools map. Using SNPEff^82^ we predicted SNP effects using the NCBI reference annotation for strain 17 and all SNPs filtered based on their DP (>=4X), MQ (>=30) and AAF ratio (>=0.8 ALT). SNPs were then filtered based on effect and only positions with predicted “MODERATE” or “HIGH” effect were further investigated, of these SNPs were adequate sequence coverage within intervals of interest were investigated. Based on their informativeness we selected a range of SNPs for further inspection (see SI Note 5). SNP effect analysis was not performed for RIJ001 due to the low sequence coverage of the strain 17 mapping.

### HSV-1 linkage disequilibrium and population genetics analysis

HSV-1 is known to be highly recombinogenic, with recombination acting to decorrelate allele frequencies with characteristic increase with physical distance. To test for the presence of recombination in the core genome alignment we first thinned core SNPs to exclude those within 80 base pairs of each other. We applied TomaHawk (https://github.com/mklarqvist/tomahawk) to estimate the pairwise r^2^ between the remaining 812 variant sites; identifying a significant decrease with genomic distance (*p*=5.73e-8). To partition the core genetic diversity into clusters we applied ADMIXTURE v1.3.0^83^, first pruning the dataset for sites in high LD (*--indep-pairwise 100 60 0.3*). ADMIXTURE was applied to the pruned alignment of 1,231 core SNPs in unsupervised mode for *K* ranging from 1-15, with the lowest cross-validation error obtained at *K*=4. Finally, a haplotype sharing analysis was applied to recover fine-scale population structure. Chromopainter v2^42^ was run in haploid mode (*-j*) in an all-versus-all manner to ‘paint’ each HSV-1 relative to all others in the dataset assuming a uniform recombination map. Chromopainter was initially run with Expectation-Maximisation (EM) to estimate the average switch rate (*-n*) and emission (*-M*) probabilities resulting in mean estimates of n=723.72 and M=0.013 which were fixed in a final run across all individuals. fineSTRUCTURE^42^ was then applied to cluster individuals based on patterns of haplotype sharing using an estimated normalisation parameter *c=*0.075 running the MCMC with 1000000 burn-in iterations (*-x*), 2000000 sampling iterations (*-y*), retaining every 10000th sample (*-z*) and applying 1000000 comparisons during the tree-building step (*-t*).

### HSV-1 phylogenetics analysis and recombination filtering

A maximum likelihood phylogenetic tree was built over the core genome alignment in IQTree v1.6.12^84^ specifying a GTR substitution model run for 1000 boot-strap iterations. To identify sites derived from recent recombination events within the core alignment we applied 3Seq^85^ to identify recombination events between all triplet pairs (Supplementary Table 4, Extended Data Fig. 3, Supplementary Note 3). All identified recombinant blocks were subsequently excluded from the alignment and a maximum likelihood phylogenetic tree constructed on the recombination filtered alignment comprising a total of 2814 SNPs as before. Phylogenies were visualised and plotted using ggTree v2.4.1^86^. To assess the possibility of recombination between our ancient genomes and HSV-2, we analysed a full genome alignment, generated using muscle^87^, made up of two HSV-1 genomes per phylogroup (KM222720, LT594457, MH999845, HM585512, HM585501, HM585496), a recombinant HSV-1 strain (HM585509), HSV-2 strains (JN561323.2, KR135308) and our ancient genomes using RDP5^45^. Using RDP/GENECOV/Chimera/MaxChi/BootScan/SiScan/3Sec we could not detect any evidence of recombination between the ancient HSV-1 strains and modern HSV-2 strains (see SI). Additionally, we used the query versus reference detection method with our ancient genomes set as reference with the same result.

### Phylogenetic Dating

We tested for the presence of a significant temporal signal over the recombination filtered core phylogeny, subset to include only those HSV-1 genomes with associated collection dates, using PhyloStems^88^ and BactDating^89^. Where a range in sampling date was provided in the associated metadata we set the date of sampling to the midpoint. While we obtained no global temporal signal basal to the tree, we obtained a significant regression following date randomization over the ancestral node common to Phylogroup I and Phylogroup II (Fig. 4) comprising all Eurasian HSV-1 strains. BEAST v2^46^ was applied to the subset alignment, specifying the constant site weights. First, bModelTest^90^ was applied to identify the site model and associated substitution model, with 74.4% of posterior support for a transversion model (TVM) (Fig. 4A). Specifying TVM as prior three possible demographic models (‘Coalescent constant’, ‘Coalescent exponential’, ‘Coalescent Bayesian Skyline’) were then run in BEAST2 with both a strict and a relaxed prior on the clock, each time specifying an MCMC chain length of 200 million sampling every 5000th from the run. In each case convergence was assessed through evaluation of the Effective Sample Size (ESS), requiring a value >200, and manual inspection of the MCMC convergence in Tracer v1.7.1. In each case models were run ‘without data’ by selecting sampling from the prior in the Beauti GUI. Finally, to assess model support, each run was repeated with nested sampling^91^ to generate a marginal likelihood for all possible model comparisons. Mean, higher posterior density estimates and posterior distributions are provided in Extended Data Table 4 and Fig. 4B. In addition, an analysis was conducted over the global phylogeny, specifying a coalescent skyline demographic model and allowing a uniform prior on substitution rates bounded by the estimates obtained for sampled Eurasian diversity (Fig. 3). For this analysis the tree prior was fixed to the maximum likelihood phylogeny estimated over the core, recombination filtered alignment. Maximum clade credibility (MCC) trees were generated using TreeAnnotator v2.6.3, discarding the first 10% of posterior trees as burn-in.

**Fig. 4.**
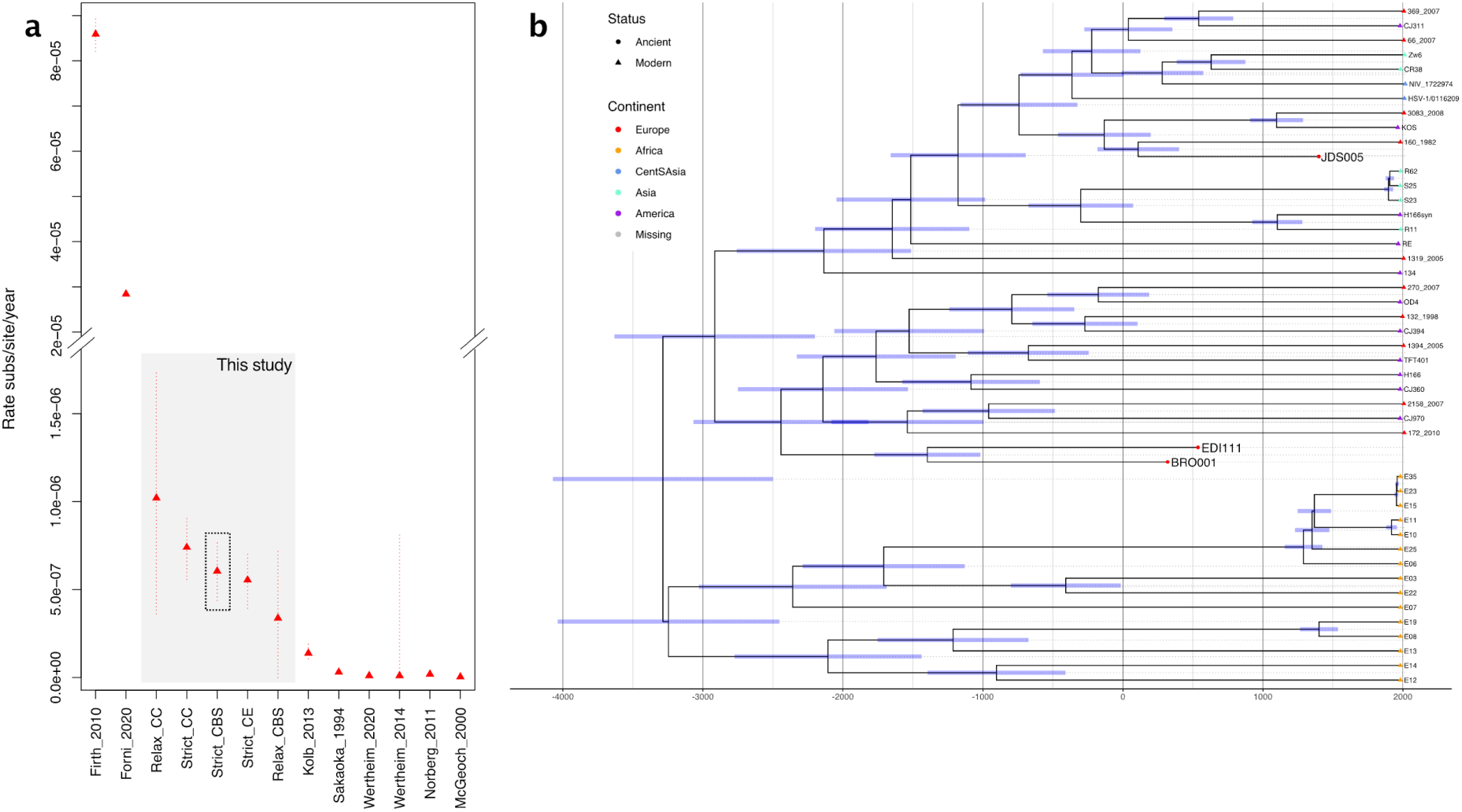
Mutation rates estimated in previous studies of HSV-1 and close relatives and time calibrated maximum clade credibility tree over a recombination pruned core genome alignment of all HSV-1 genomes. **a,** Y-axis provides the rates recovered measured in substitutions per site per year. X-axis documents eight published studies including five estimates obtained in this study using ancient genomes. Firth et al 2010 and Forni et al 2020 provide tip-calibration estimates assuming a power law rate decay model. Other studies make various assumptions about co-divergence: Kolb et al 2013 (assumption of human demographic history eg. split of 34 thousand between Europe/North America), Sakaoka et al 1994 (assuming co-divergence with host), Wertheim et al 2020 (assumption of codivergence within the McHV-1 viruses and their corresponding five Macaca host species), Wertheim et al 2014 (calibrated assuming viral-host codivergence eg. tMRCA of old and new world primates), Norberg et al 2011 (BEAST applied to US7 and US8, divergence with HSV-2 set to 8.45 M years) and McGeoch et al 2000 (assuming co-divergence with host). Our most likely model estimate is highlighted in grey with estimates in this study, making use of ancient calibration points, falling between those estimated from modern diversity assuming a power law rate decay and those estimated assuming a scenario of co-divergence. Confidence intervals are indicated by a dotted line. **b,** The time calibrated phylogeny is inferred following specification of a strict clock model and coalescent skyline population priors assuming a uniform prior bounded by the rates estimated for the Eurasian branch exhibiting temporal signal (Extended Data Fig. 4A,B). Modern samples are represented by triangular tip-points, ancient samples are depicted with circular tip-points. Blue bars provide the 95% HPD interval of the estimated age at each node. Continental origins of the genomes are denoted by the tip-color as given in the legend at right.

### Mapping and quality control of human data

Raw data was returned in the form of single and paired FASTQ files. Sequences of adaptors and indexes and poly-G tails occurring due to the specifics of the NextSeq500/550 and Hiseq4000 technology were removed from the ends of sequences using cutadapt v1.9^63^ and AdapterRemoval v2.0^92^. Sequences shorter than 30 bp were removed to avoid random mapping of sequences from other species.

Sequence reads were mapped to the human reference sequence (GRCh37/hg19) using Burrows-Wheeler Aligner (BWA v0.7.12) ^93^ command aln with seeding disabled. After mapping, the sequences were converted to BAM format and only sequences that mapped to the reference genome were retained using samtools v1.9^69^. Next, multiple bams from the same individual, but different runs were merged using samtools merge. Reads with mapping quality under 30 were filtered out and duplicates were removed with picard v2.12 (http://broadinstitute.github.io/picard/index.html). MapDamage2.0^72^ was applied to estimate the frequency of 5’ C > T transitions. Values are reported in Supplementary Table 1A.

We estimated contamination using the mitochondrial DNA using a method from^94^, which aligns the raw mtDNA reads to the RSRS^95^, determines the haplotype using GATK pileup^96^, counts the number of heterozygous reads on haplotype-defining sites as well as adjacent sites and calculates a ratio that takes into account ancient DNA damage by excluding positions where a major allele is C or G and the minor is T or A respectively. Secondly we performed a similar calculation on the X chromosome in JDS005, a male individual. This method was not applied for EDI111, a female, nor for BRO001 as the Y chromosome coverage was low (Extended Data Table 2). Samtools-1.9^69^ stats was used to determine the number of final reads, average read length, average coverage etc.

### Genome-wide *H. sapiens* analysis and population genetics

Genetic sex was calculated using the script sexing.py from^97^, estimating the fraction of reads with mapping quality > 30 mapping to the Y chromosome out of all reads mapping to either the X or Y chromosome. Genetic sexing confirmed morphological sex estimates.

Due to differences in genomic coverage between the samples and generally lower than optimal coverage (15X) for calling heterozygous variants, the genotypes were estimated in three ways: with GATK-3.5 -T HaplotypeCaller^71, 96^ using the default settings; with ANGSD-0.916^98^ command --doHaploCall, sampling a random base for all genomic positions and pseudo-haploidised by copying the sampled allele; and using an imputation pipeline detailed in Hui et al.^99^. VCF files were annotated with the ClinVar database downloaded on 27/01/2020 and filtered for clinical significance “pathogenic”, “likely pathogenic” and for “herpes” information tags (Supplementary Table 10).

We calculated runs of homozygosity using hapROH^100^ (v0.3a3) (Extended Data Table 3). Following the recommended setting, we used pseudo-haploidised genotypes at polymorphic sites in the 1000 Genomes Project panel as the input and kept all parameters at their default values (roh_in=1, roh_out=20, roh_jump=300, e_rate=0.01, e_rate_ref=0.0, cutoff_post=0.999, max_gap=0, roh_min_l=0.01). When summarising results for each individual, we also followed the default settings for filtering and merging ROH segments (snp_cm=50, gap=0.5, min_len1=2.0, min_len2=4.0).

## Extended Data Legends

**Extended Data Table 1 | Archaeological summary**

Site, burial, archaeological and radiocarbon dating, skeletal age estimate, morphological and genetic sex estimate, mitochondrial and Y chromosome (where applicable) haplogroup information for each ancient individual in this study.

**Extended Data Table 2 | Contamination estimates**

Results of mitochondrial and X chromosome-based estimates of human contamination for each sample in this study.

**Extended Data Table 3 | Runs of homozygosity in samples over 1x coverage**

List of runs of homozygosity segments in JDS005 and EDI111 as estimated from imputed genomes by hapROH^100^.

**Extended Data Table 4 | BEAST2 posterior estimates**

BEAST2 posterior estimates following assessment of the results over three demographic models and tested using both strict and relaxed clock priors. Marginal likelihood and SD estimates are also provided. The analysis highlighted in italics failed to converge following 200 million iterations.

**Extended Data Fig. 1 | Phylogenetic networks with different reference genomes**

SplitsTree neighbour nets for core genome alignments obtained when using three distinct reference genomes **a,** 2158_2007, **b,** S25, **c,** E07. Ancient samples are highlighted with a diamond with accessions names as per Supplementary Table 5.

**Extended Data Fig. 2 | LD vs. genetic distance, CV for ADMIXTURE and Recombination regions**

**a,** Decline in LD (r 2, y-axis) over genetic distance (x-axis) specifying a minimum r2 value of 0.25 to aid visibility. Following linear regression *R*^2^ 0.1, *p*-value 5.73x10^-8^. Confidence interval indicated by grey shaded area. **b,** ADMIXTURE cross-validation estimates following application of unsupervised clustering. Inferred ancestry components at K=4 are provided in Figure 2d. **c,** Putatively recombinant sites identified by 3Seq. Blue provides individual recombination blocks identified in triplet parent child combinations (see Supplementary Table 6). Red panel at top provides the combined recombinant tracts excluded over the HSV-1 reference genome.

**Extended Data Fig. 3 | Maximum Likelihood Phylogeny**

Maximum likelihood phylogenetic tree over the HSV-1 dataset. Tip colours provide the continent of sampling, with the three ancient samples shown with circles and all other tips (modern samples) having triangular symbols. Panel at right provides the phylogroup assignment for those samples overlapping with Pfaff et al. 2016 as denoted in the legend. Green *s denote nodes with >70% boot-strap support following 1000 boot-strap iterations of the tree building step.

**Extended Data Fig. 4 | Testing for temporal signal, BModelTest and Posterior Densities**

**a,** Linear relationship between the time of sample collection (x-axis) and root-to-tip phylogenetic distance (y-axis) estimated from a recombination pruned maximum likelihood phylogenetic tree over the global phylogenetic dataset. Confidence intervals are indicated by dashed lines. **b,** Following identification of a significant temporal signal at the node falling basal to the non-African HSV-1, the same plot is demonstrated only for descendants from this node (as highlighted in panel a). In both cases plot headings provide the rate, MRCA, regression coefficient and *p*-value following 10,000 permutations of the sampling date computed using the BactDating roottotip() function. **c,** Models with blue circles are inside 95%HPD, red outside, and without circles have at most 0.40% support. Model 123421 (transversion model – TVM) had 86.79% posterior support and 74.40% cumulative support. **d,** Posterior distributions estimated under three possible specifications of demographic models using a strict molecular clock. **e,** Posterior distributions estimated under three possible specifications of demographic models using a relaxed molecular clock. In both panel **d** and **e** red - coalescent constant model, green - coalescent exponential model, blue - coalescent bayesian skyline model.

### Availability of data and material

The ancient datasets generated and analysed during the current study are available in the ENA repository under the accession ID: PRJEB46097. Accession IDs for modern comparative data used are listed in Supplementary Table 5. All other data supporting the findings of this study are available within the paper and its supplementary information files or available from the corresponding author upon reasonable request.

## Acknowledgements

We thank the support of the Cambridge Archaeological Unit, Quinton Carroll and colleagues at Cambridgeshire County Council as well as other members of the After the Plague project: Toomas Kivisild, Piers D. Mitchell, Bram Mulder, Tamsin O’Connell and Jay T. Stock. We also acknowledge Mary Price for her isotopic analysis of JDS005. We also would like to acknowledge archaeologist Elizaveta M. Chernykh and physical anthropologist Ivan G. Shirobokov. This work is supported by the Wellcome Trust (Award no. 2000368/Z/15/Z) and St John’s College, Cambridge (J.E.R., S.A.I, C.C., A.R., C.L.S.); The Max Planck Society (J.K., A.H.); the Estonian Research Council grant PUT (PRG243) (L.S., A.S., M.M., C.L.S) and PUT (PRG1027) (K.T., L.S., M.G., A.K.); and the European Union through the European Regional Development Fund (Project No. 2014-2020.4.01.16-0030) (C.L.S., M.G., M.M.); the European Regional Development Fund (Project No. 2014-2020.4.01.15-0012) (M.M.); and the ERC Synergy Grant HistoGenes (No. 856453) (A.H.). L.vD is supported by a UCL Excellence Fellowship. C.J.H acknowledges support from the NIHR Cambridge Biomedical Research Centre Antimicrobial Resistance theme. A.R acknowledges support from the British Archaeological Association.

## Author Information

### Authors’ contributions

C.L.S., J.E.R., J.K., and A.H. conceived the study. S.A.I., C.C., A.K., L.V., and E.M.C. assembled skeletal samples and S.A.I. and J.D. performed osteological analysis. C.C. performed dating analyses. A.R. generated and analysed isotopic data. C.L.S, L.S., K.T., and A.S. generated ancient genome shotgun data. M.G. designed and implemented the viral target capture. J.K. and A.H. provided additional shotgun sequencing and expertise. M.M. and J.E.R. provided funding. L.vD., M.G., C.L.S., C.J.H., E.d’A. and R.H. analysed the genomic data. C.J.H. provided expertise on herpes viruses. C.L.S. wrote the manuscript with the input of all co-authors.

## Ethics declarations

### Competing interests

The authors declare no competing interests.

## Notes

### Competing Interest Statement

The authors have declared no competing interest.

