## Supplementary Notes for "Ancient herpes simplex 1 genomes reveal recent viral structure in Eurasia"

### Note 1. Archaeological sites, material, and radiocarbon determination

*Craig Cessford, Sarah Inskip, Jenna Dittmar and Aivar Kriiska*

**The Hospital of St. John, Cambridge, UK**

The Hospital of St. John, Cambridge was a medieval establishment for the care of the poor and infirm, with the exception of pregnant women, lepers, the wounded, crippled and insane [1], the assemblage of 400 internments (*c*. 1204-1511)[2] included a young adult male (JDS005 / PSN031[1] / Burial: 198; Skeleton: 1232 ) aged 17-25 who almost certainly died between 1350-1450 CE. JDS005 is a young adult male individual that has very little evidence of pathology on the skeleton (75-100% complete). There is no evidence of endocranial lesions. Nor is there any evidence of new bone formation on the axial skeleton. This individual presents with both active and healed cribra orbitalia, which may be related to hematopoietic marrow conversion related to anemia[2]. There is evidence of dental enamel hypoplasia present on the teeth, which suggest that this individual experienced a 'disruption' in the body's ability to form dental enamel during childhood. This is caused by periods of physiological stress (illness, malnutrition etc.). On the tibiae, there is striated lamellar bone (evidence of a healed periosteal reaction, which is the formation of new bone in response to injury or other stimuli of the periosteum surrounding the bone). This has various possible causes: trauma (unlikely in this case), cancer (unlikely), or things like chronic irritation of the area, or a non-specific infection (more likely).

The skeletal evidence indicates poor dental health. This individual had minor to moderate periodontal disease and most teeth have minor to moderate calculus present along the

[1] PSN numbers are project numbers from the After the Plague project.
[2] This remains controversial in the field

cemento-enamel junction, with the roots of the anterior teeth also affected. A gross carious lesion (which has resulted in destruction of more than 50% of the tooth) is present on the right second maxillary molar. This alveolus has been destroyed by a large abscess that has perforated through the buccal aspect of the right maxilla (affecting M1 and M2). The left second mandibular molar was lost antemortem. The alveolus is very well healed, which indicates that this tooth was lost at least several months prior to the death of this individual. This individual also experienced chronic maxillary sinusitis. It is possible that this could be related to the observed dental pathology but this is unlikely (because the maxillary sinus was not perforated by the abscess).

Although this individual lived to adulthood, the presence of these 'non-specific stress indicators' suggests that he experienced ill-health early on. These 'indicators' have been linked to increased risk of mortality.

**Barrington A, Edix Hill, Cambridgeshire, UK**

The site of Barrington A, Edix Hill, Cambridgeshire is an early Ango-Saxon cemetery in use between 500 - 650 CE. EDI111 (PSN650 / grave 38, skeleton 127A), an adult female (35-45 years old), with evidence of arthritis, dental disease and trauma was interred in a double burial, the other inhumation is a 1% complete sub-adult (2-3 years old)[3]. The lower extremities of the skeleton of 127A were badly damaged by ploughing activity[3]. The burial was originally assigned to the early phase of the cemetery (500-575 AD), although this phasing is based solely on the presence of amber beads in the burial which is relatively weak evidence. The radiocarbon determination supports but does not categorically confirm this phasing (Supplementary Table 8).

Information on the radiocarbon determination for EDI111 is given in Supplementary Table 8.

EDI111 has been radiocarbon dated to 545–590 calCE (1σ interval) or 437–636 calCE (12σ interval). As a lower left 2nd molar was sampled, the radiocarbon determination does not date the death of the individual but when the tooth formed by age 11 - 13. As this individual was osteologically assessed as 35–45 years old when they died, this suggests that the individual died 20 - 30 years after the intervals quoted. Given the date and location of the Edix Hill cemetery it is unlikely that the individuals buried there consumed marine fish, and the δ13C value of -20.4 is compatible with this. Low level consumption of freshwater fish by those buried at Edix Hill is possible, but can not be accurately assessed. Consumption of freshwater fish would have the effect of making the radiocarbon determination older than the death of the individual. The impact of the potential freshwater reservoir effect has not been estimated and is likely to be slight.

**Brodovsky, Nevolino, Russia**

The Brodovsky burial site is located in the village of Borody in Perm Krai of Russia. On the old riverbank of Shakva several barrows and ground burials have been found[4]. Archaeological excavations have been carried out there repeatedly since 1898[4,5]. By the burial customs and grave goods, the Brodovsky burial site is related to the Nevolino culture, which existed in Prikamaye from 4$^{th}$ to 9$^{th}$ centuries CE[5,6]. Unfortunately, the analysed tooth comes from a burial, which can no longer be more precisely loca-lized.

**Raadhuisstraat 185-193, Alphen aan den Rijn, South-Holland, NL**

The individuals buried on the right bank of the river Rhine are most likely victims of a French attack on the villages Bodegraven and Zwammerdam in december 1672[3]. The inhabitants of the villages were massacred and some of the bodies ended up in the Rhine

[3] Palstra, S.W.L., Raadhuisstraat Cold Case (2020),

which took them down stream. Some bodies were washed ashore on the excavated area[4], which was still unoccupied at that time[5]. A total amount of four complete individuals were discovered. The bodies were all men from European descent. Two bodies (S36 and S37) were buried disrespectfully together in one grave and face down, which suggests they were French soldiers (i.e. the enemy).

Individual S16 was a male of 26-35 years old[6]. In his youth he probably suffered from an insufficiency of vitamin D, which resulted in a slight bent in the lower legs and a deviation in the ribs. The man was a fervent smoker of clay pipes. Traces of the habit are visible in multiple places on the teeth, where the hard clay pipe, usually put in the same place in the mouth, has worn the teeth (see fig.).

---

[4] Koekkelkoren, A.M.H.C. *et al.* Raadhuisstraat 185-193, Alphen aan den Rijn, Gemeente Alphen aan den Rijn, *IDDS Archeologie rapport* **2313** (2021)

[5] Vitters, M. Stille getuigen van het rampjaar 1672 (2021)

[6] Veselka, B. Menselijk skeletmateriaal afkomstig uit de Raadhuisstraat 185-193, Alphen aan den Rijn Fysisch Antropologische Rapportage (2019)

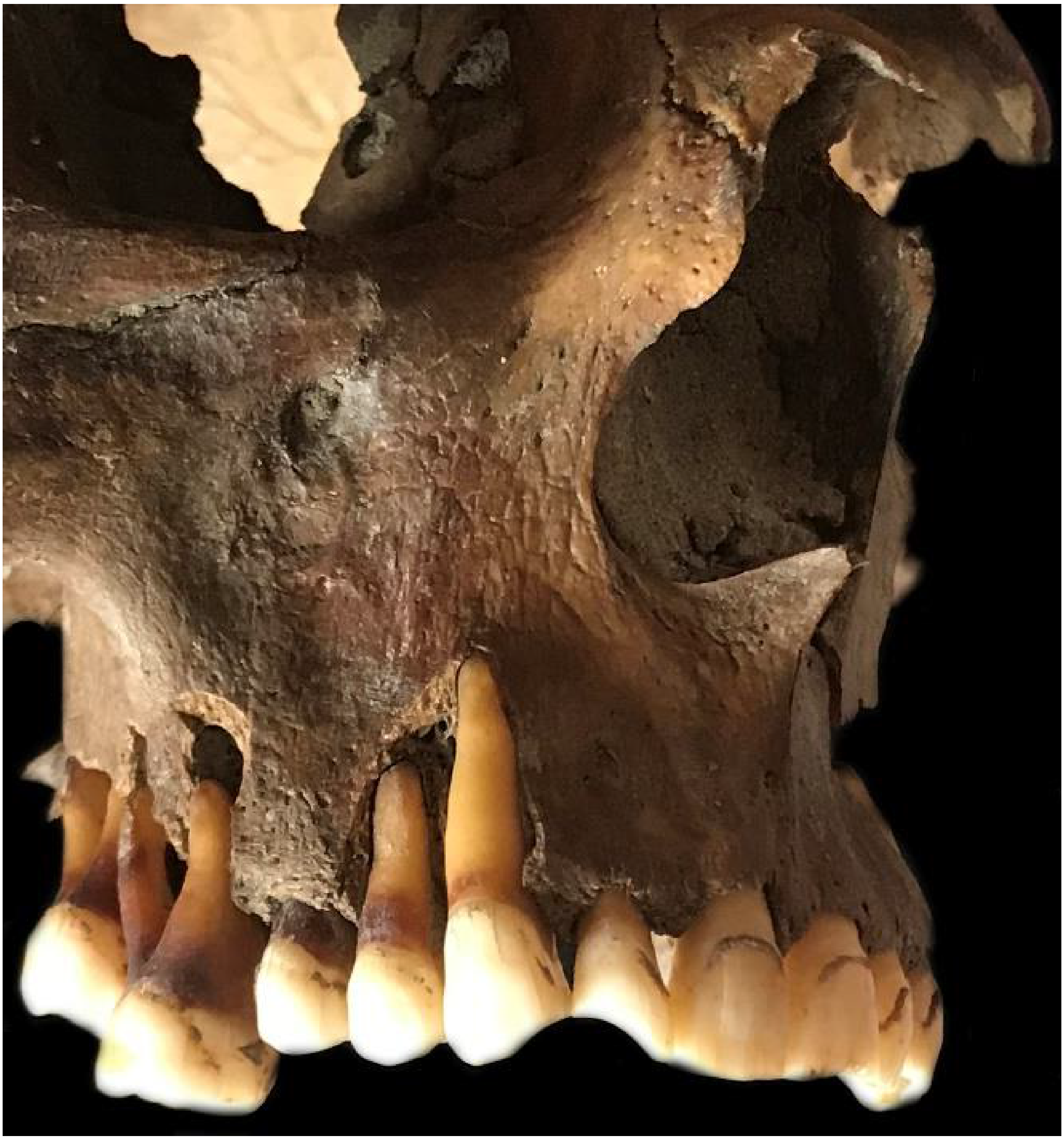

#### Note 2. Generation of ancient DNA

*Christiana Scheib and Meriam Guellil*

##### Sampling

Teeth were sampled from skeletons using gloves. Molars were preferred due to having more roots and larger mass, but premolars were also sampled.

##### Generation of aDNA libraries

Sampling, decontamination, extraction, purification and library preparation for JDS005 was carried out as described in Scheib et al. 2018[7]. EDI111 was processed at the Institute of Genomics ancient DNA facility at the University of Tartu as described in Scheib et al. 2019[8]. BRO001 was processed at the Institute of Ecology and Earth Sciences ancient DNA facility at the University of Tartu as described in Saag et al. 2019[9].

##### Sequencing

The JDS005 sample was initially shotgun-sequenced on the Illumina NextSeq500/550 using the High-Output single-end 75 cycle kit at the DNA Sequencing Facility in the Department of Biochemistry, University of Cambridge. Once HSV-1 reads were identified in the library using KRAKEN[10], the library was sent to the Max Planck Institute for the Science of Human History, Jena, Germany for eleven additional lanes of sequencing on HiSeq4000 paired-end 150 cycle kit. Since the libraries were single-indexed and pooled with post-capture libraries for *M. leprae, S. enterica* and *Y. pestis*, these lanes cannot be used for additional metagenomic searches due to cross-contamination/index-hopping from the capture libraries.

EDI111 was shotgun sequenced on the Illumina NextSeq500/550 using the High-Output single-end 75 cycle kit at the University of Tartu Institute of Genomics Core Facility in three separate runs, the first multiplexed with other samples, the second two alone followed by three additional runs at the DNA Sequencing Facility in the Department of Biochemistry, University of Cambridge.

BRO001 was shotgun sequenced on the Illumina NextSeq500/550 using the High-Output single-end 75 cycle kit at the University of Tartu Institute of Genomics Core Facility in a multiplex with other samples. Once HSV-1 reads were identified, the sample was subjected to a custom designed multi-viral capture as described below.

**Metagenomic screening**

Generated shotgun sequencing data was inspected with fastqc[11], and quality filtered and trimmed of adaptors using cutadapt[12] with the settings -m 30 --nextseq-trim=20 for single-end data with multi-core support enabled (-m 30 --nextseq-trim=20 --times 3 -e 0.2 -j 0 --trim-n). We then deduplicated the filtered FASTQ files using ParDre[13]. Following these steps, microbial DNA was assigned to taxa using KrakenUniq[14]. The custom database contained complete genomes and chromosome level assemblies of bacteria, viruses, archaea and protozoa. The human genome, the NCBI Viral Neighbor database and the contaminant databases UniVec and EmVec were also included. A custom E-value was calculated as follows: $(\frac{K}{R}) \times C$. Here, K is the *k*mer count, R is the read count and C is the coverage of the taxon *k*mer dictionary. The E-value cut-off for further inspection was 0.001.

**Capture**

Due to low coverage, we enriched the shotgun library of sample BRO001 and RIJ001 for HSV-1 DNA using an Arbor Biosciences Custom MyBaits multispecies viral capture kit (v4), that includes sequences from 283 *Herpes simplex virus type 1* genome sequences and 261 *Herpes simplex virus type 2* genome sequences. The captured library was amplified using 2X KAPA HiFi HotStart ReadyMix DNA Polymerase and primers IS5 and IS6[15]. Following amplification, the library was sequenced on a NextSeq500 platform (150bp, paired-end) at the Estonian Biocentre.

#### Note 3. Genome-wide analysis of Human herpes simplex virus 1

*Meriam Guellil, Lucy van Dorp and Christiana L. Scheib*

##### Alignment to the reference sequence

Raw data was returned in the form of single and paired FASTQ files. Prior to mapping single-end datasets were trimmed and filtered as described above for the metagenomic screening. Paired-end datasets were also filtered and trimmed using cutadapt[12] (-m 30 --nextseq-trim=20 --times 3 -e 0.2 -j 0 --trim-n --pair-filter=any). Forward and reverse reads were then merged using FLASH[16] (-z 125).

All datasets were merged by library and mapped non-competitively against the *Herpes simplex type 1* reference sequences for strain 17 (NC_001806.2), strain S25 (HM585513.2), strain 2158_2007 (LT594106.1) and strain E07 (HM585497.2), to exclude a reference bias, using bwa aln (-n 0.1 -l 1000) and bwa samse[17]. We then used samtools[18] to convert and sort the data to bam format. Duplicates were marked and removed using Picards's[19] MarkDuplicates module. We then merged sequencing runs for all libraries using samtools merge (-R) and realigned reads around indels using GATK's[20] RealignerTargetCreator and IndelRealigner modules. We estimated deamination rates using mapDamage2.0[21]. Alignments were visualized using a python script [22] using matplotlib[23]and mapping statistics were collected using Qualimap2[24]. Edit distances were computed with a custom python script [25] using the modules numpy[26], pysam[27] and pysamstats[28]. Additionally, we also mapped our data non-competitively to the *Herpes simplex type 2* strain HG52 reference genome (NC_001798.2) using the same workflow.

All positions were stored using samtools v1.10 mpileup (*-aa* flag)[18]. Variant positions were then called using bcftools v1.10.2, filtering for sites with a mapping quality of at least 20, a depth of 3, a quality of 20 and taking forward those positions with an alternate allele fraction >0.8 (MQ>20, DP>3, Q>20, AAF>0.8). Consensus fasta files were generated using bcftools *consensus* masking those regions of the genome not covered over the reference.

We estimated the mappability across the reference sequence strain 17 using GenMap[29] (-K30 -E2). Coverage across gene intervals was calculated using bedtools coverage[30] and mean mappability across these intervals was computed using bedtools map. Using SNPEff[31] we predicted SNP effects using the NCBI reference annotation for strain 17. SNP with HIGH or MODERATE impact predictions were inspected manually based on filtering criteria (see Note 5).

**Genotyping**

We additionally performed a genotyping analysis on our alignments to the reference sequence HSV-1 strain 17. Grouping of HSV-1 into three distinct genotypes (A, B and C) is commonly based on intragenic SNPs in the glycoprotein genes *US4* (*gG*) and *US7* (*gI*) [32] and the more recently gene *US2*[33]. Due to low coverage at some genotyping coordinates, not all SNPs could be assessed in the ancient samples. This led to ambiguous genotypes in gene *US2* and *US4*. Interestingly, most genotyping loci were covered in our RIJ001 alignment, which was too low coverage for phylogenetic analysis. The sample can probably be typed as (B,C)/B/B. Genotypes are detailed in Supplementary Table 11. However, consistent with previous findings[34], the genotyping scheme does not seem to be correlated with the phylogenetic clustering.

**Supplementary Table 11 | Results of genotyping of HSV-1 strains included in this study**

| Gene/Publication | SNP | JDS005 | | EDI111 | | BRO001 | |
|---|---|---|---|---|---|---|---|
| | | Base | Group | Base | Group | Base | Group |
| US2 (Glück et al. 2015) | 134171-5 | GGGAT | A/B | GGGAT | A/B | GGGAC | C |
| | 134185 | A/G* | A/B/C | A | A | G | A/B/C |
| | 134217 | *N/A* | - | A | A | G | B/C |
| | 134237 | ??G* | A | *N/A* | - | CCT | B/C |
| | 134241 | A* | A | *N/A* | - | G | B/C |
| US4/gG (Norberg et al. 2006) | 137088 | *N/A* | - | *N/A* | - | *N/A* | - |
| | 137090 | *N/A* | - | *N/A* | - | *N/A* | - |
| | 136974 | C* | A/B | T* | C | C | A/B |
| US7/gI (Norberg et al. 2006) | 140160 | G | A/B | G | A/B | *N/A* | - |
| | 140298 | T | A | G* | B/C | G | B/C |
| Result | | | A/(A/B)/A | | A/C/B? | | C/(A/B)/(B/C) |

* Only 1X coverage at this position

#### HSV-1 linkage disequilibrium and population genetics analysis

HSV-1 is known to be highly recombinogenic, with recombination acting to decorrelate allele frequencies with characteristic increase with physical distance. To test for the presence of recombination in the core genome alignment we first thinned core SNPs to exclude those within 80 base pairs of each other. We then applied TomaHawk (https://github.com/mklarqvist/tomahawk) to estimate the pairwise $r^2$ between the remaining 812 variant sites; identifying a significant decrease in $r^2$ with genomic distance ($p$=5.73e-8). To partition the core genetic diversity into clusters we applied ADMIXTURE v1.3.0[35], first pruning the dataset for sites in high LD (*--indep-pairwise 100 60 0.3*) inlight of the presence of recombination within the dataset. ADMIXTURE was applied to the resulting alignment encompassing 1,231 core SNPs in unsupervised mode for values of $K$ ranging from 1-15, with the lowest cross-validation error obtained at $K$=4. Finally, a haplotype sharing analysis was applied to recover fine-scale patterns of population structure. Chromopainter v2[36] was run in haploid mode (*-j*) in an all-versus-all manner to 'paint' each HSV-1 relative to all others in the dataset. A uniform recombination map was applied assuming a rate of 0.00001 per base pair. Chromopainter was initially run employing Expectation-Maximisation (EM) to estimate the average switch rate (*-n*) and emission (*-M*) probabilities resulting in mean estimates of n=723.72 and M=0.013 which were fixed in a final run across all individuals (*-a 0 0*). fineSTRUCTURE[36] was then applied to cluster individuals based on patterns of haplotype sharing using an estimated normalisation parameter $c$=0.075 running the MCMC with 1000000 burn-in iterations (*-x*), 2000000 sampling iterations (*-y*), retaining every 10000th sample (*-z*) and applying 1000000 tree comparisons during the tree-building step (*-t*).

**Compilation of comparative HSV data**

All publicly available assemblies of HSV-1 were downloaded from NCBI by querying NCBIVirus using TaxID 10298 pre-specifying a genome length between 150000 to 200000. In addition, all accessions not captured by these filters but included in Forni et al.[37] were included resulting in a dataset of 299 modern HSV-1 samples (Supplementary Table 5). Within this dataset, accessions were filtered to include only single sample representatives (from studies employing longitudinal sampling) and to exclude heavily passaged strains or characterised recombinants (see notes in Supplementary Table 5). The metadata was manually curated to add country and date information through consultation of the dataset prepared by Forni et al.[37] and through specific requests to associated research groups.

Our interest is in whether regions of the HSV-1 genome can be used for phylogeographic inference thus two main points were considered in choosing comparative samples: geographic origin and mutation rate. The 'natural' mutation rate of HSV-1 in immunocompetent human hosts is lower than in cell culture or immunocompromised individuals[38], thus strain from these conditions or studies were left out along with strains sequenced for purposes other than clinical study such as using novel synthetic sequencing methods. Strains isolated from large clinical settings in which the likely geographic origin of the host/strain is not noted were also discarded. An initial analysis considered 162 genomes (including the reference Strain17) which were subsequently filtered to remove those deriving from cosmopolitan centers, for instance excluding genomes sampled in the USA post-dating 1989 which otherwise dominated the dataset. A total of 64 genome sequences were retained for phylogenetic analysis (Supplementary Table 5).

**Preparation of genome sequences**

Variants were called for all included modern genomes from their assemblies using the snippy pipeline and specifying contigs as input (https://github.com/tseemann/snippy). All modern and ancient samples were then jointly considered to construct a core genome alignment using snippy-core, as in Lassalle et al[39], considering variant positions observed at least ten times across the dataset and masking difficult to call positions including repeat genes (RL1, RL2, RS1) together with all regions identified by Szpara et al. 2014[40] relative to reference strain 17. This resulted in a core SNP alignment including 3,815 well defined variant positions of which each of our ancient genomes (BRO001, EDI111, JDS005) had 275, 185 and 286 core SNPs respectively. Equivalent analyses were conducted using three representative alternate references to assess the placement of the ancient strains.

**HSV-1 phylogenetics analysis and recombination filtering**

A maximum likelihood phylogenetic tree was built over the core genome alignment in IQTree v1.6.12[41] specifying a GTR substitution model run for 1000 boot-strap iterations. To identify sites derived from recent recombination events within the core alignment we applied 3Seq[42] to identify recombination events between all triplet pairs (Supplementary Table 6, Extended Data Fig. 2C). All identified recombinant blocks were subsequently excluded from the alignment and a maximum likelihood phylogenetic tree constructed on the recombination filtered alignment in IQTree as before. Phylogenies were visualised and plotted using ggTree v2.4.1[43]. To assess the possibility of recombination between our ancient genomes and HSV-2 strains, we analysed a full genome alignment, generated using muscle[44], made up of two representative HSV-1 genomes per phylogroup (NC_001806.2, LT594457, MH999845, HM585512, HM585501, HM585496), a recombinant HSV-1 strain (HM585509) , HSV-2 strains (JN561323.2, KR135308) and our ancient genomes (except RIJ001) using RDP5[45]. Our analysis using RDP/GENECOV/Chimera/MaxChi/BootScan/SiScan/3Sec could not detect any recombination event between our ancient genomes and the HSV-2 strains. Additionally we used the query versus reference detection method with our ancient genomes set as reference with the same result.

**Phylogenetic Dating**

We tested for the presence of a significant temporal signal over the recombination filtered core phylogeny, subset to include only those HSV-1 genomes with associated collection dates, using PhyloStems[46] and BactDating[47]. Where a range in sampling date was provided in the associated metadata we set the date of sampling to the midpoint. While we obtained no global temporal signal basal to the tree, we obtained a significant regression over the ancestral node common to Phylogroup I and Phylogroup II (Extended Data Fig. 4A,B) comprising all Eurasian HSV-1 strains. Bayesian tip-dating was applied to this subset SNP alignment, specifying the constant site weights, using BEAST v2[48]. First, bModelTest[49] was applied to estimate the best support site model and associated substitution model, with 74.4% of posterior support for a transversion model (TVM) (Extended Data Fig. 4C). Specifying TVM as prior three possible demographic models (‘Coalescent constant’, ‘Coalescent exponential’, ‘Coalescent Bayesian Skyline’) were then run in BEAST2 both specifying a strict prior on the clock rate and a relaxed prior on the clock, each time specifying an MCMC chain length of 200 million sampling every 5000th from the run. In each case convergence was assessed through evaluation of the Effective Sample Size (ESS) requiring a value >200 and manual inspection of the MCMC convergence in Tracer v1.7.1. In each case models were run ‘without data’ by selecting sampling from the prior in the Beauti GUI. Finally, to assess model support, each run was repeated with nested sampling[50] to generate a marginal likelihood and Bayes factor support for all possible model comparisons. Mean, higher posterior density estimates and posterior distributions are provided in Extended Data Table 2 and Extended Data Fig. 4D. In addition, an analysis was conducted over the global phylogenetic diversity, specifying a coalescent skyline demographic model and allowing a uniform prior on substitution rates bounded by the estimates obtained for the node common to phylogroup I and phylogroup II. For this analysis the tree prior was fixed to the maximum likelihood phylogeny estimated over the core, recombination filtered alignment. In all cases the maximum clade credibility (MCC) trees were generated using TreeAnnotator v2.6.3, discarding the first 10% of posterior trees as burn-in.

#### Note 4. Genome-wide analysis of *H. sapiens*

*Christiana Scheib*

##### Alignment to the reference sequence and quality control

The sequence reads were mapped to the human reference sequence (GRCh37/hg19) using Burrows-Wheeler Aligner (BWA 0.7.12)[17] command aln with seeding disabled. After mapping, the sequences were converted to BAM format and only sequences that mapped to the reference genome were kept using samtools 1.9[18]. Next, multiple bams from the same individual, but different runs were merged using samtools merge, reads with mapping quality under 30 were filtered out and duplicates were removed with picard 2.12 (http://broadinstitute.github.io/picard/index.html).

##### aDNA Authentication

Samtools-1.9[18] option stats and BAMstats-1.25 (http://bamstats.sourceforge.net/) were used to determine the number of final reads, average read length, average coverage etc. As a result of degradation over time, aDNA can be distinguished from modern DNA by certain characteristics: short fragments and a high frequency of C > T substitutions at the 5' ends of sequences due to cytosine deamination. The program mapDamage2.0[21] was used to estimate the frequency of 5' C > T transitions. JDS005 had an average of 20% C > T frequency in the 5' ends and EDI111 12.5%.

To estimate the level of potential contamination in the libraries (Extended Data Table 2), we used the human genome in two ways, the first estimating mtDNA contamination was estimated using the method from[51], which aligns the raw mtDNA reads to the RSRS[52], determines the haplotype using GATK pileup[53] counts the number of heterozygous reads on haplotype-defining sites as well as adjacent sites and calculates a ratio that takes into account ancient DNA damage by excluding positions where a major allele is C or G and the minor is T or A respectively. The second performed a similar calculation on the X chromosome in JDS005, a male individual. This second method was not used for EDI111, a female.

**Genetic sex estimation**

Genetic sex was calculated using the script https://github.com/pontussk/ry_compute from Skoglund et al. 2013[54], estimating the fraction of reads with mapping quality > 30 mapping to Y chromosome out of all reads mapping to either X or Y chromosome. Genetic sexing confirmed morphological sex estimates.

**Determining mtDNA haplogroups**

Raw reads were mapped to the revised Cambridge Reference Sequence[55] and the resulting bam files were indexed using samtools-1.9[18]. Variants were called using Samtools 1.9 mpileup variant-only option[18] and filtered using bcftools v 1.1[18]. Haplogroups were assigned using Phylotree build 16[56] accessed at www.phylotree.org, Haplogrep[57] accessed at https://haplogrep.uibk.ac.at.

**Y chromosome variant calling and haplotyping**

Y chromosome variants were called in JDS005 as haploid and picking one allele at random (--doHaploCall 1) in ANGSD-0.916 [58] and filtered for regions that uniquely map to Y chromosome, retaining 8.8 Mb, when using short read sequencing technology[59]. Haplogroup assignments were made on the basis of in silico genotyping of the samples for 108,000 informative variants 1000 Genome Project populations[60] in 456 geographically diverse high-coverage Y chromosome sequences[59] and those annotated by https://isogg.org/tree/ and https://www.yfull.com/. In haplogroup labelling we followed the nomenclature of Karmin et al. (2015).

**Variant calling and imputation of genotypes**

Due to differences in genomic coverage between the samples and generally lower than optimal coverage (>15X) for calling heterozygous variants were called in three ways: GATK-3.5 -T HaplotypeCaller using the default settings, with ANGSD[58] command --doHaploCall, sampling a random base for all genomic positions and pseudo-haploidised by copying the sampled allele, and lastly by using an imputation pipeline detailed in Hui et al. 2020[61]. We estimated the genotypes with BEAGLE 4.1[62] using genotype likelihoods produced by ATLAS[63] used in Beagle -gl mode, followed by imputation in Beagle -gt mode from sites whose GP exceed 0.99. To balance between imputation times and input and accuracy, we used 606 European sliding window of 2000 SNPs and northern Han Chinese (CHB) genomes from Phase 3 of the 1000 Genomes Project as the reference panel in Beagle -gl step, and all the 2,504 genomes from the above dataset in Beagle -gt step. Only sites with GP above 0.99 and minor allele frequency above 0.3 in the EUR+CHB panel were included in phenotypic analysis.

**Phenotype Prediction**

We analysed a total of 75 variants involved in diet and immunity response. by selecting 2 Mb around each variant and merging the overlapping region, for a total of 48 regions from 17 autosomes and the X chromosome.

We call the variants using ATLAS v0.9.0 [64] task=call and method=MLE commands at positions with a minimum allele frequency (MAF) ≥ 0.1% in the UK10K panel (extracted from the HRC, Haplotype Reference Consortium panel)[65]. After calling the variants separately for each sample, we merged them in one VCF file per region. We used the VCFs as input for the first step of our imputation pipeline [61] (genotype likelihood update), performed with Beagle 4.1 -gl command [66] using the same panel as before as reference. We then discarded the variants with a genotype probability (GP) less than 0.99 and imputed the missing genotype with the -gt command of Beagle 5.0 [67] using the HRC as a reference panel. We then discarded the variants with a GP < 0.99 and used the remaining SNPs to perform the phenotype prediction. Results are reported in Supplementary Table 10.

#### Note 5. Functional analysis of ancient HSV-1

*Charlotte Houldcroft and Meriam Guellil*

**Variant annotation**

We annotated our vcf files using snpEff[31] with a custom database for the HSV-1 reference strain based on the gff3 for NC_001806.2. For the annotation, we also included indels in our vcfs, which were inspected by eye to avoid misalignments and misidentifications. Using bedtools[30] and our GenMap data we further computed the number of SNPs per gene and the coverage in gene intervals (>MQ30). Variants are listed in Supplementary Table 3.
We investigated moderate and high impact SNPs within glycoprotein intervals associated with an immune response based on the following criteria: effect grade, coverage and depth of coverage of the interval of interest, MQ>=30 at the position and DP>=3 at the position.
<u>Selected SNPs:</u>

**JDS005**

*UL27: gB* - Two SNPs predicted to be moderate but they do not fall into Uniprot-predicted features, such as the virion surface. Unlikely to be significant.

*UL22: gH* - Four non-synonymous SNPs (p.Ala150Thr; p.Ser138Ala; p.Thr127Ile; p.Pro110Ser) are all in the Uniprot-predicted virion surface portion of the glycoprotein, increasing the chances that these variants may be recognised as epitopes by the immune system[68]. Glycoprotein gH also has a role in cell infection (fusion of the virus for entry)[69].

*US4: gG* - This is a short protein, so it is hard to predict the effect of a non-synonymous SNP at aa3.

*US6: gD* - This SNP is not in the transmembrane domain of gD, but lies just outside the N terminus (amino acids 1-29), which are recognised; speculated to be neutral[70].

**EDI111**

*UL27: gB* - One moderate effect SNP, but AA873 is an intravirion (predicted) domain, so less likely to be an epitope.

**BRO001**

*US3*: - One high effect (stop_loss) SNP in this region, which is a significant virulence factor for HSV-1. This region encodes serine/threonine kinase.

*UL27: gB* - One moderate effect SNP. As in EDI111, it lies within an intravirion (predicted) domain, so less likely to be an epitope.

*UL22: gH* - A number of predicted moderate SNPs are in the Uniprot-predicted virion surface portion of the glycoprotein, increasing the chances that these variants may be recognised as epitopes by the immune system.

*US4: gG* - This is a short protein, so it is hard to predict the effect of a non-synonymous SNP at aa3 (same as the JDS genome, thus it may be a common polymorphism).

*US6: gD* - Just outside the predicted transmembrane domain of gD; speculated to be neutral.

#### Note 6. Host Susceptibility analyses

The ancient human genomes were called as detailed in Supplementary Note 4 and vcf files were annotated with the latest ClinVar vcf downloaded from the ftp site on 27/01/2020. The resulting annotated snps were filtered for clinical significance "pathogenic", "likely pathogenic" and for "herpes" information tags (Supplementary Table 9). For JDS005 (~11X) variants with DP ≤ 3 were discarded; however, since EDI111 had a much lower coverage of ~6X, all variants with DP ≥ 2 were kept and evaluated by hand. Variants called homozygous were accepted. Due to aDNA damage present and average genomic coverage below 15X, variants called heterozygous were assessed as either likely heterozygous (ratio of ref:alt at ~1:1) or likely erroneous (ratio of ref:alt >2:1 and a C > T or G > A transition). Interestingly, though not related to HSV-1, JDS005 carries an autosomal dominant variant for nocturnal frontal lobe epilepsy (rs12721510, Supplementary Table 9), which may explain why he was in the Hospital of St John the Evangelist at a young age.

In addition, specific genomic regions of interest for susceptibility to Herpes viruses were identified by searching ClinVar for "Herpes" and including all genes in which pathogenic SNPS were listed. This included 14 genes. A bed file of gene positions was used to extract the haplotypes from a merged VCF file using bcftools [71].

Herpesviridae have been found to be more abundant in cases of chronic and aggressive periodontitis[72,73] though the connection with HSV-1 remains inconclusive[74,75]. Sequencing reads from EDI111 point to the presence of bacteria commonly associated with minor to moderate periodontal disease along with a number of other microorganisms that may indicate poor dental health.

#### Note 7. Isotope Analysis

Analysis of carbon and nitrogen isotopes in archaeological human dentine and bone collagen can be used to provide information about diet in the past[76,77]. This is based on the principle that carbon and nitrogen isotope values in skeletal tissues largely reflect the isotopic composition of the diet consumed during life[78,79] and these signatures are preserved in archaeological skeletal remains. Broadly, carbon isotope values can provide information about the types of plants consumed ($C_3$ vs $C_4$) [80,81], nitrogen isotope values can provide information about animal protein consumption and subsequent trophic level[76,82,83] and analysis of a combination of carbon and nitrogen isotope values can provide information about marine fish consumption[82,84,85]. In addition, analysing multiple skeletal tissues from an individual can provide information about diet across an individual's lifespan[86]. As primary dentine does not remodel[87], analysis of carbon and nitrogen isotopes in dentine is representative of diet at the time of formation in childhood, whereas bone constantly remodels throughout life, at varying rates[88] and therefore isotope values in bone collagen represent diet in the years before death[76].

**Generation of isotopic data**

Dentine collagen from the root (enamel-dentine junction to root apex) of an upper left 2$^{nd}$ molar (representing diet from approx.8.5-12.5yr[89] from JDS005 was analysed, and combined with collagen isotope data generated by Price (2013)[90] from a rib midshaft (representing diet in the years before death[88]). EDI111 and BRO001 were not analysed as they were outside the scope of the After the Plague project.

Preparation of collagen samples was carried out in the Dorothy Garrod Laboratory, McDonald Institute, University of Cambridge, following the laboratory standard operating procedures based on a modified version of the Longin method (Longin 1971, Collins and

Galley 1998, Richards and Hedges 1999). An approx. 300mg sample of root dentine and a 500-1000mg sample of rib bone were cut using a hand-held Dremel drill with a diamond-tipped cutting wheel. The sample surfaces were then abraded using a sandblaster to remove surface contaminants. To extract the collagen, samples were first demineralized in approximately 8ml of cold 0.5M aq. hydrochloric acid (HCl), then rinsed with distilled water and heated in approximately 8ml of pH 3.0 H2O for 48hrs at 75°C until gelatinized. The samples were then filtered using Ezee filters (60-90μm), frozen (-20°C, then -80°C) and placed into a freeze-drier until fully lyophilized. Analysis of the collagen samples was carried out in the Godwin Laboratory, Department of Earth Sciences, Cambridge. All samples were analysed in triplicate. For each one of the triplicates, 0.8mg (±0.1mg) of lyophilised collagen was weighed into a tin capsule. For each batch of samples submitted for analysis, a suite of in-house standards were also weighed into tin capsules and analysed (caffeine, alanine, nylon, protein, EMC). Analysis was carried out using a Costech automated elemental analyser coupled with a Thermo Finnigan MAT253 isotope ratio mass spectrometer in continuous flow mode. All samples are reported on the international scale relative to VPDB for carbon and AIR for nitrogen (Hoefs 1997). Based on replicate analyses of standards, analytical error was <±0.2‰. All results were quality checked to see if they were within the acceptable atomic C:N ratio range of 2.9-3.6 (DeNiro 1985) and above the minimum acceptable %C and %N threshold ( >4-5% for C and >13% for N (Ambrose 1990)).

**Results**

Both tissues from JDS005 provided successful $\delta^{13}C$ and $\delta^{15}N$ values: dentine collagen $\delta^{13}C$ = -19.5‰ and $\delta^{15}N$ = 9.9‰, rib collagen $\delta^{13}C$ = -19.5‰ and $\delta^{15}N$ = 9.8‰. The C:N ratios were within the acceptable range of 2.9-3.6[91] and the %C and %N in the samples were above the minimum thresholds (taken as >13% for C and >4-5% for N[92].

The isotope values for JDS005 are below the mean for the general adult population from the Hospital of St John (mean dentine collagen $\delta^{13}C$ = -19.2‰, $\delta^{15}N$ = 12.1‰, n=55; mean rib collagen $\delta^{13}C$ = -19.0‰, $\delta^{15}N$ = 12.5‰, n=114)[93]. More specifically, they are below the mean for adult males in the Hospital (mean dentine collagen $\delta^{13}C$ = -19.1‰, $\delta^{15}N$ = 12.1‰, n=31; mean rib collagen $\delta^{13}C$ = -18.9‰, $\delta^{15}N$ = 12.6‰, n=64) and general Parish adult males from the nearby High and Late Medieval site of All Saints by the Castle (mean dentine collagen $\delta^{13}C$ = -19.5‰, $\delta^{15}N$ = 11.9‰, n=14; mean rib collagen $\delta^{13}C$ = -19.3‰, $\delta^{15}N$ = 12.5‰, n=25) (ibid.). Indeed, JDS005 has some of the lowest $\delta^{15}N$ values observed out of all the High and Late Medieval adults sampled from the Cambridgeshire population (ibid.).

JDS005 is also unusual in the lack of meaningful change in the isotope values between their dentine and rib collagen (difference in values is below the analytical error of ±0.2‰). This is different to the rest of the adult population from the Hospital of St John, who generally exhibited a change in values between the tissues, particularly in $\delta^{15}N$ (mean $\Delta^{13}C_{rib\ coll\text{-}dentine\ coll}$ = 0.1‰, mean $\Delta^{15}N_{rib\ coll\text{-}dentine\ coll}$ = 0.5‰, n=53). JDS005 is well below the mean $\Delta^{15}N_{rib\ coll\text{-}dentine\ coll}$ values for adult males in the Hospital (0.7‰, n=30).

Overall, the isotope values for JDS005 are indicative of a $C_3$ plant based diet, with little regular terrestrial animal protein input and little to no marine protein input. The lack of change in isotope values between the dentine and rib collagen indicates no detectable change in diet between childhood and adulthood. In terms of $\delta^{15}N$ values, JDS005 appears to be quite unusual compared to the rest of the population buried in the Hospital of St John, as well as the Cambridge town population as a whole. Their much lower values and lack of change could indicate they were consistently consuming less animal proteins than much of the rest of the population of the town. When taken in context with their young age at death, pathologies present, short stature and eventual burial in the Hospital, this may be taken as an indication of someone who could not afford regular animal or marine proteins in their diet, someone who was too sickly to consume them, or even someone who chose not to consume them.

#### Note 8. History of Kissing

*Christiana Scheib*

While there are Neolithic figurines that have been interpreted as a couple embracing[94], the earliest known record of kissing is a Bronze Age manuscript from South Asia[95] (Fig. 4B). The custom may have made its way back to the Mediterranean with the return of Alexander the Great's troops around 300 BCE. Three hundred years later, the Emperor Tiberius is said to have tried to ban kissing at official functions in an effort to stop the spread of disease (unclear whether the disease was herpes)[96].

Such a transition in transmission routes would allow for transmission within larger host communities as well as at the intersection of populations when they meet. Sexual-romantic kissing is far from universal in human cultures (and is also observed in Chimpanzees and Bonobos), but is higher in prevalence in modern European, Middle Eastern and Asian cultures[97].

27. *pysam*. (Github).

28. Miles, A. *pysamstats*. (Github).

39. Lassalle, F. *et al.* Whole genome sequencing of Herpes Simplex Virus 1 directly from human cerebrospinal fluid reveals selective constraints in neurotropic viruses. *Virus Evol* **6**, veaa012

(2020).

87. Gage, J. P. *Collagen and dental matrices*. (Wright, 1989).

88. Fahy, G. E., Deter, C., Pitfield, R., Miszkiewicz, J. J. & Mahoney, P. Bone deep: Variation in stable isotope ratios and histomorphometric measurements of bone remodelling within adult

humans. *J. Archaeol. Sci.* **87**, 10–16 (2017).

89. AlQahtani, S. J. *Atlas of Human Tooth Development and Eruption*. (2008).
90. Price, M. How Did the Social Changes of the Fourteenth Century Affect the Everyday Lives of English People? A Case Study from St. John's Divinity School, Cambridge. (University of Cambridge, 2013).
91. DeNiro, M. J. Postmortem preservation and alteration of in vivo bone collagen isotope ratios in relation to palaeodietary reconstruction. *Nature* **317**, 806–809 (1985).
92. Ambrose, S. H. Preparation and characterization of bone and tooth collagen for isotopic analysis. *J. Archaeol. Sci.* **17**, 431–451 (1990).
93. Rose, A. Life in Medieval Cambridge: an isotopic analysis of diet and mobility. (University of Cambridge , 2020).
94. Malone, C. & Stoddart, S. Figurines of Malta. in *The Oxford Handbook of Prehistoric Figurines* (ed. Insoll, T.) (Oxford University Press, 2017).
95. Parkinson, R. B. *The Tale of Sinuhe and Other Ancient Egyptian Poems, 1940-1640 BC*. (Oxford University Press, 1998).
96. Danesi, M. *The History of the Kiss!: The Birth of Popular Culture*. (Palgrave Macmillan, New York, 2013).
97. Jankowiak, W. R., Volsche, S. L. & Garcia, J. R. Is the romantic-sexual kiss a near human universal? *Am. Anthropol.* **117**, 535–539 (2015).
