## Extended Data for "Ancient herpes simplex 1 genomes reveal recent viral structure in Eurasia"

### Extended Data Items

#### Extended Data Table 1 | Archaeological summary

**Extended Data Table 1 | Archaeological, anthropological and genetic summary of the individuals used in this study**

| Sample ID | Site | Burial No. | Skeleton ID | Sample taken | Morphological age estimate | Chronological dating | Morphological sex estimate | Genetic sex estimate | Mitochondrial Haplogroup | Y chromosome Haplogroup |
|---|---|---|---|---|---|---|---|---|---|---|
| JDS005 | Hospital of St John | 2710 | 1232 | LLM1 | Young Adult (17-25) | 1350-1450 CE* | Male | XY | T1a1 | J1a2 |
| EDI111 | Barrington A, Edix Hill | 38 | 127A | LLM2 | Adult (35-45) | 1520 ± 27 BP | Female | XX | H1b | N/A |
| BRO001 | Nevolino | n/a | n/a | PM2 | Adult | 1680 ± 35 BP | n/a | XY | U4a1d | R1b |
| RIJ001 | Alphen aan den Rijn | 11 | 16 | ULM1 | | 17th century* | | | | |

Radiocarbon dates are uncalibrated and * indiciates that the sample dating is inferred from site stratigraphy utilising radiocarbon dates of in situ material.

#### Extended Data Table 2 | Contamination estimates

**Extended Data Table 2 | Summary of contamination estimates from human genomes**

| Sample ID | Run | mtDNA based contamination | X chromosome based contamination | SNP sites | SNP sites with flanking | Method1: old | | Method1: new | | Method2: old | | Method2: new | |
|---|---|---|---|---|---|---|---|---|---|---|---|---|---|
| | | | | | | MoM | ML | MoM | ML | MoM | ML | MoM | ML |
| JDS005 | Merged | 0.21% | 0.69% | 54,748 | 492,732 | 0.7763% ± 4.8e-4 | 0.6023% ± 6.5e-14 | 0.7705% ± 4.8e-4 | 0.6027% ± 7.0e-13 | 0.7721% ± 1.1e-3 | 0.6284% ± 1.3e-13 | 0.7663% ± 1.1e-3 | 0.6280% ± 1.7e-12 |
| EDI111 | Merged | 0.17% | N/A | | | | | | | | | | |
| BRO001 | Run1 | 0.06% | N/A | | | | | | | | | | |
| RIJ001 | Run1 | 1.05% | N/A | | | | | | | | | | |

X-based contamination estimates can only be run on males with sufficient autosomal coverage, in this case only JDS005 fit the criteria.

#### Extended Data Table 3 | Runs of homozygosity

**Extended Data Table 4 | Runs of homozygosity in samples over 1x genomic coverage**

| Sample ID | max_roh | sum_roh>0.1 | n_roh>0.1 | sum_roh>0.5 | n_roh>0.5 | sum_roh>1 | n_roh>1 | sum_roh>2 | n_roh>2 | sum_roh>4 | n_roh>4 | sum_roh>8 | n_roh>8 |
|---|---|---|---|---|---|---|---|---|---|---|---|---|---|
| JDS005 | 2.15 | 70.68 | 55 | 70.68 | 55 | 70.68 | 55 | 4.21 | 2 | 0.00 | 0 | 0.00 | 0 |
| EDI111 | 2.82 | 77.16 | 59 | 77.16 | 59 | 77.16 | 59 | 4.87 | 2 | 0.00 | 0 | 0.00 | 0 |

#### Extended Data Table 4 | BEAST2 posterior estimates

**Extended Data Table 3 | BEAST2 posterior estimates following assessment of the results over three demographic models and tested using both strict and relaxed clock priors**

| Demographic model | Posterior_ESS | Likelihood | TreeHeight | ClockRate | Marginal_Likelihood | SD |
|---|---|---|---|---|---|---|
| **Strict Clock** | | | | | | |
| Coalescent_constant | 24303.305 | -191833.726(-191843.326--191825.36) | 6138.328(4844.956-7830.772) | 7.400e-07(5.568e-07-9.160e-07) | -192077.699 | 18.39 |
| Coalescent_exponential | 2109.163 | -191834.328(-191843.986--191825.872) | 8063.489(5982.904-10795.248) | 5.542e-07(3.930e-07-7.179e-07) | -192127.163 | 18.20 |
| Coalescent_Bayes_Sky | 22405.288 | -191795.129(-191804.854--191786.913) | 7268.874(5477.789-9591.929) | 6.052e-07(4.380e-07-7.773e-07) | -192048.638 | 16.93 |
| **Relaxed Clock** | | | | | | |
| Coalescent_constant | 2507.823 | -191627.308(-191639.602--191616.991) | 4634.172(2524.156-9330.682) | 1.028e-06(3.611e-07-1.752e-06) | -193976.069 | 45 |
| Coalescent_exponential | *82.523* | *-191788.646(-191800.533--191777.469)* | *9903.537(3846.677-22354.934)* | *4.560e-07(1.4589e-07-9.361e-07)* | */* | */* |
| Coalescent_Bayes_Sky | 3243.329 | -191642.994(-191654.969--191632.129) | 13834.335(3992.156-102241.329) | 3.3691e-07(4.373e-11-7.322e-07) | -193603.489 | 41 |

Note: Marginal likelihood and SD estimates are also provided. The analysis highlighted in italics failed to converge following 200 million iterations.

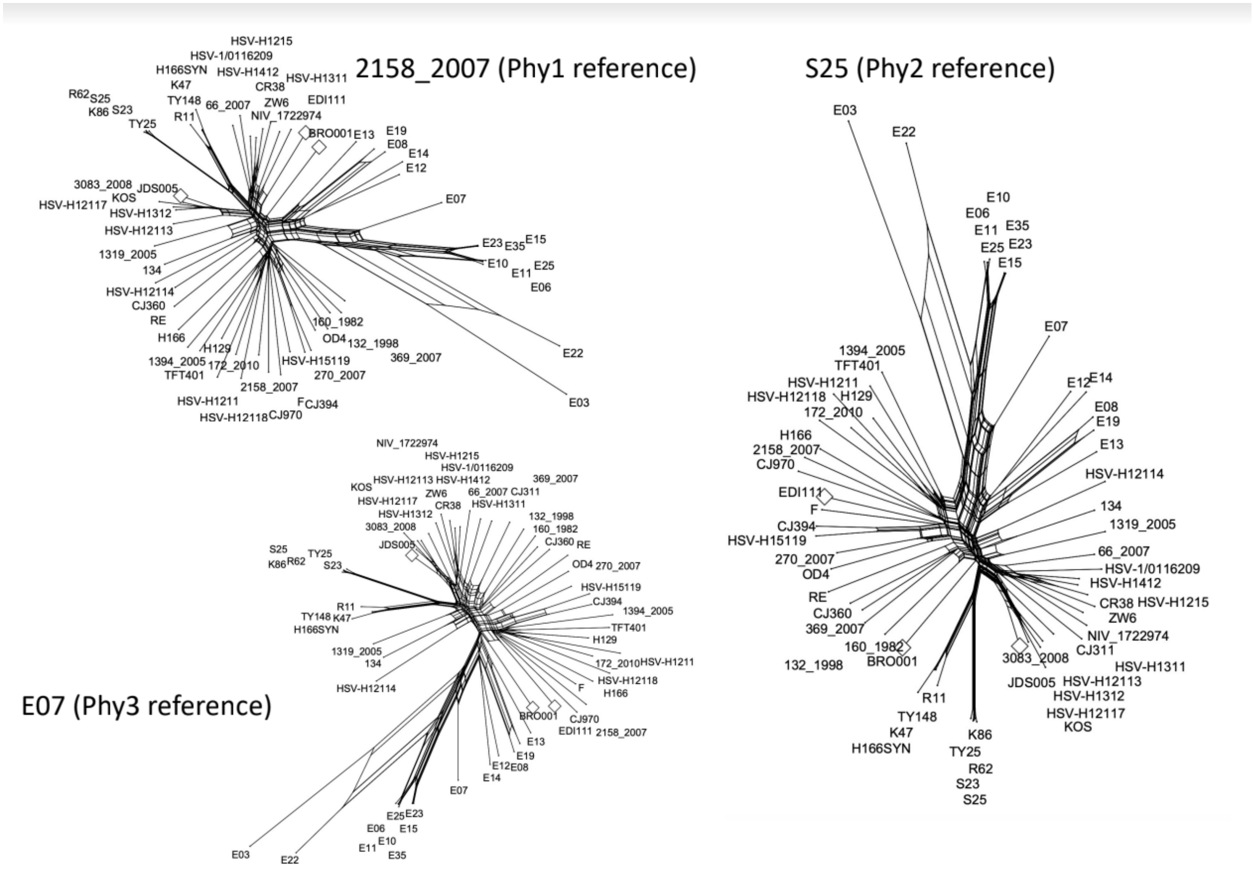


**Extended Data Fig. 1 | Phylogenetic networks with different reference genomes**

SplitsTree neighbour nets for core genome alignments obtained when using three distinct reference genomes **a,** 2158_2007, **b,** S25, **c,** E07. Ancient samples are highlighted with a diamond with accessions names as per Supplementary Table 5.

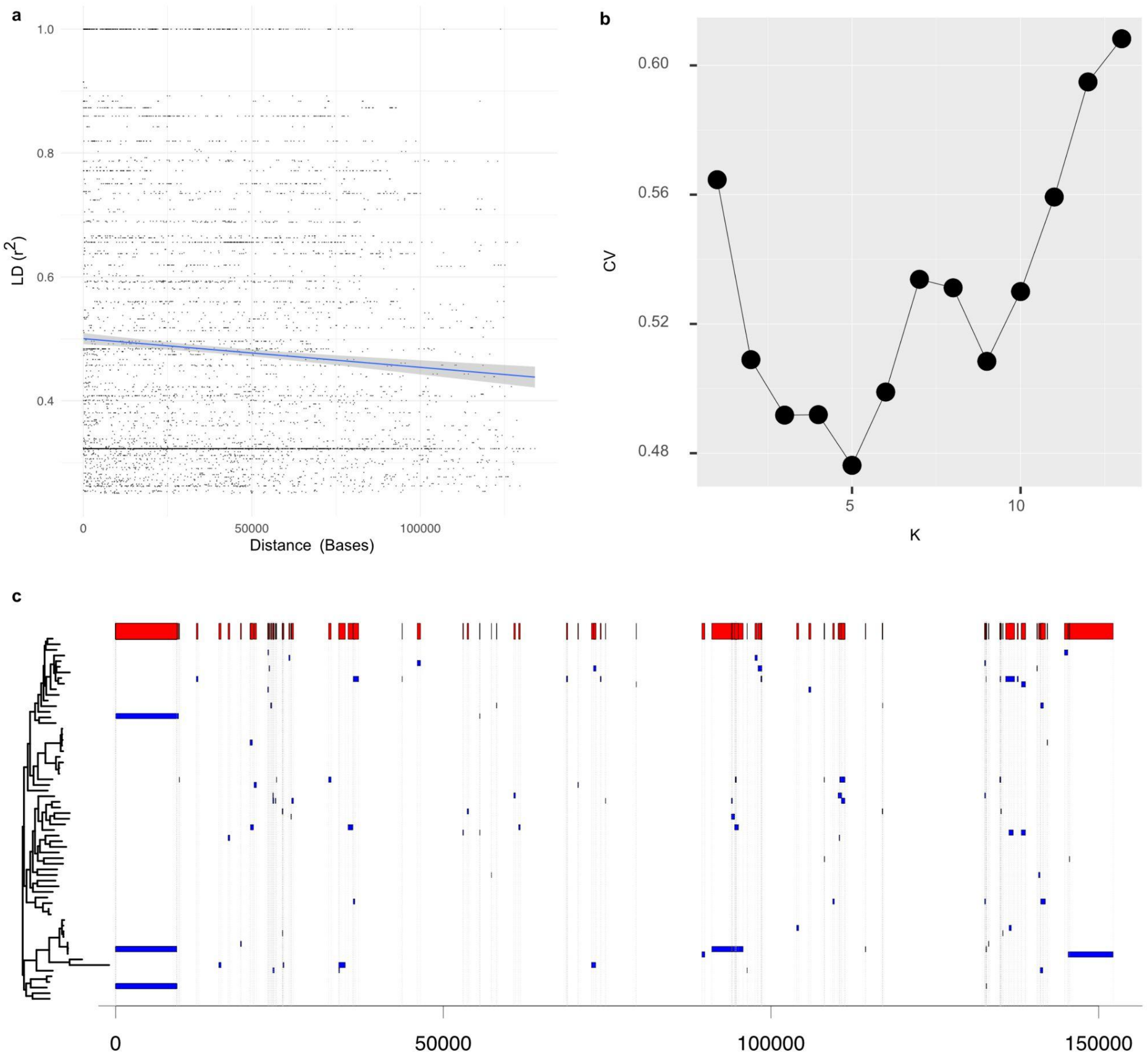


**Extended Data Fig. 2 | LD vs. genetic distance, CV for ADMIXTURE and Recombination regions**

**a,** Decline in LD (r 2 , y-axis) over genetic distance (x-axis) specifying a minimum r2 value of 0.25. to aid visibility. Following linear regression $R^2$ 0.1, p-value 0.004. Confidence interval indicated by grey shaded area. **b,** ADMIXTURE cross-validation estimates following application of unsupervised clustering. Inferred ancestry components at K=5 are provided in Figure 3c. **c,** Putatively recombinant sites identified by 3Seq. Blue provides individual recombination blocks identified in triplet parent child combinations (see Supplementary Table 6). Red panel at top provides the combined recombinant tracts identified over the HSV-1 reference genome.

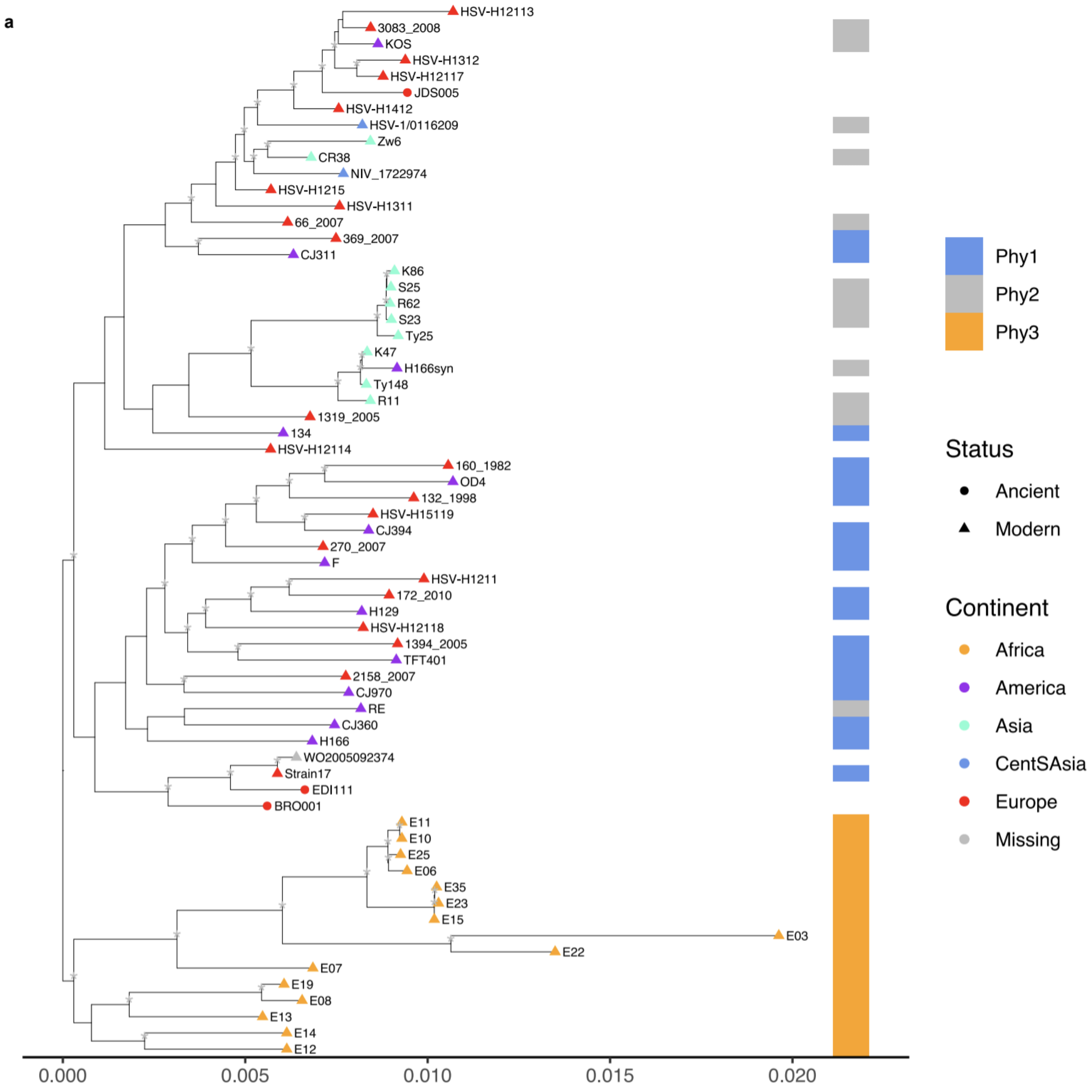


**Extended Data Fig. 3 | Maximum Likelihood Phylogeny**

Maximum likelihood phylogenetic tree over the HSV-1 dataset. Tip colours provide the continent of sampling, with the three ancient samples shown with circles and all other tips (modern samples) having triangular symbols. Panel at right provides the phylogroup assignment for those samples overlapping with Pfaff *et al.* 2016 as denoted in the legend. * denote nodes with >70% boot-strap support following 1000 boot-strap iterations of the tree building step.

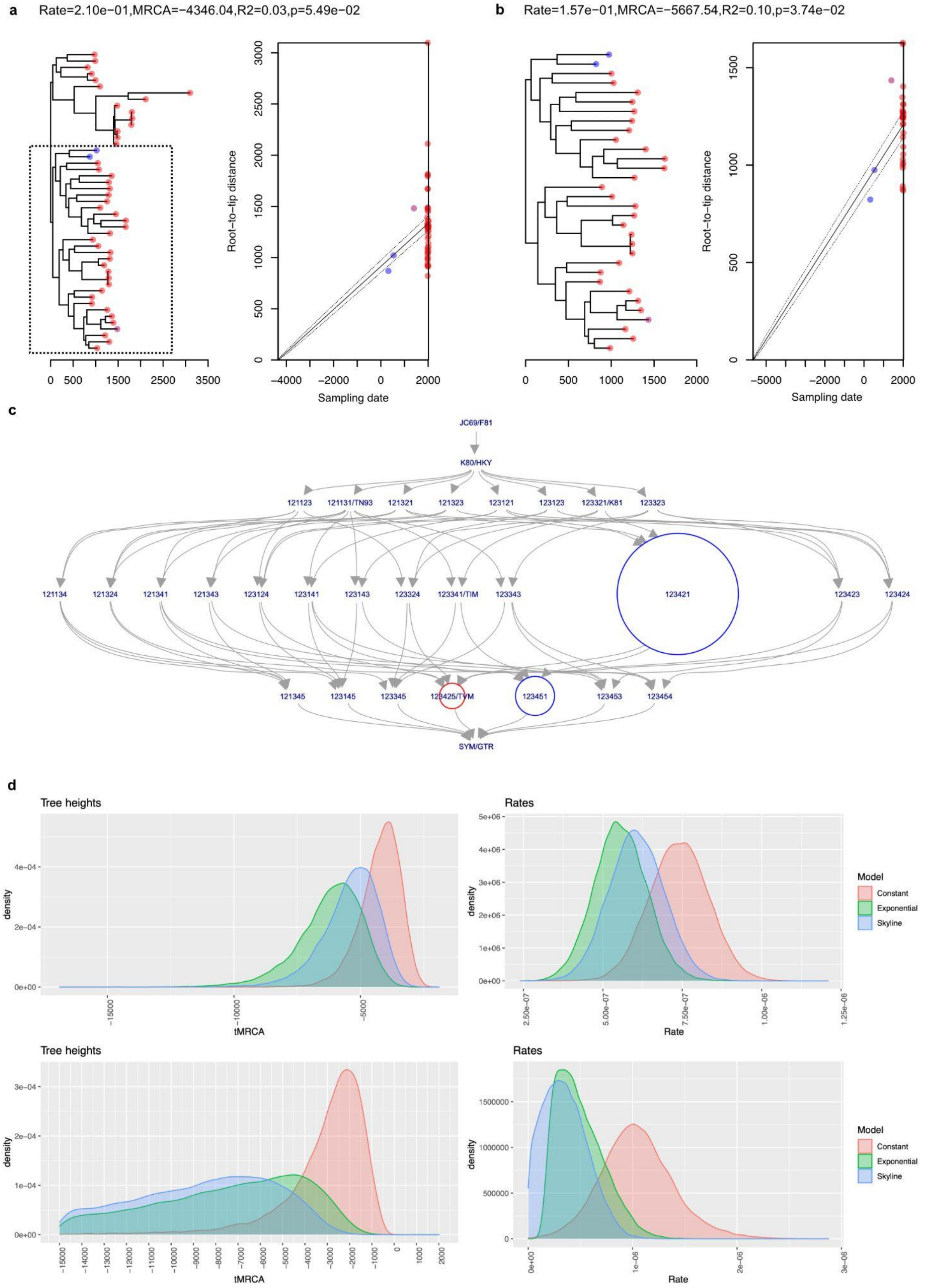
a
Rate=2.10e−01,MRCA=−4346.04,R2=0.03,p=5.49e−02
0 500 1500 2500 3500
Root−to−tip distance
0 500 1000 1500 2000 2500 3000
−4000 −2000 0 2000
Sampling date
b
Rate=1.57e−01,MRCA=−5667.54,R2=0.10,p=3.74e−02
0 500 1000 1500 2000
Root−to−tip distance
0 500 1000 1500
−5000 −2000 0 2000
Sampling date
c
JC69/F81
K80/HKY
121123
121131/TN93
121321
121323
123121
123123
123321/K81
123323
121134
121324
121341
121343
123124
123141
123143
123324
123341/TIM
123343
123421
123423
123424
121345
123145
123345
123425/TVM
123451
123453
123454
SYM/GTR
d
Tree heights
density
tMRCA
Rates
density
Rate
Model
Constant
Exponential
Skyline
Tree heights
density
tMRCA
Rates
density
Rate
Model
Constant
Exponential
Skyline

**Extended Data Fig. 4 | Testing for temporal signal, BModelTest and Posterior Densities**

**a,** Linear relationship between the time of sample collection (x-axis) and root-to-tip phylogenetic distance (y-axis) estimated from a recombination pruned maximum likelihood phylogenetic tree over the global phylogenetic dataset. Confidence intervals are indicated by dashed lines. **b,** Following identification of a significant temporal signal at the node falling basal to the non-African HSV-1, the same plot is demonstrated only for descendants from this node (as highlighted in panel a. In both cases plot headings provide the rate, MRCA, regression coefficient and p-value following 10,000 permutations of the sampling date computed using the BactDating roottotip() function. **c,** Models with blue circles are inside 95% HPD, red outside, and without circles have at most 0.40% support. Model 123421 (transversion model – TVM) had 86.79% posterior support and 86.79% cumulative support. **d,** Posterior distributions estimated under four possible specifications of demographic models (colours as per legend). Estimated tree heights are given in the left-hand panels, estimated clock rates are provided in the right-hand panels.

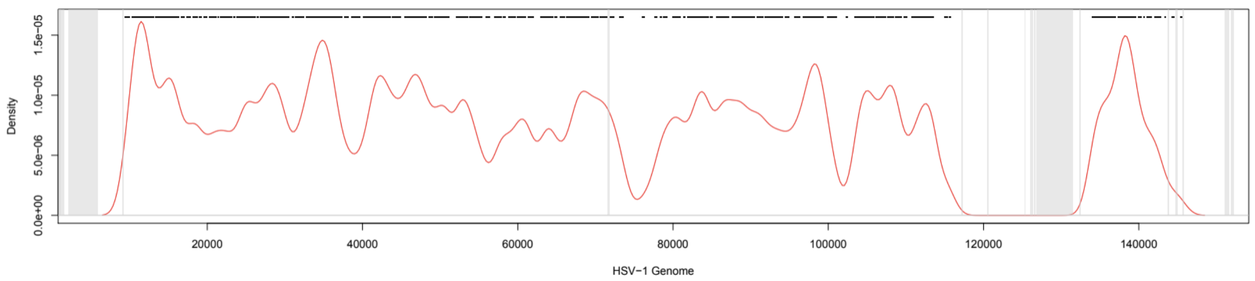


**Extended Data Fig. 5 | Distribution of variant positions called in the core genome alignment over the HSV-1 genome (x-axis).** Grey bars provide the regions masked from the alignment. The red line provides the density of variants (y-axis) with individual SNPs (n=3815) additionally demarcated by the black points atop of the figure.
